# Metabolic profiling of adiponectin levels in adults: Mendelian randomization analysis

**DOI:** 10.1101/126789

**Authors:** Borges Maria Carolina, Barros Aluísio JD, Ferreira Diana L Santos, Casas Juan Pablo, Horta Bernardo Lessa, Kivimaki Mika, Kumari Meena, Usha Menon, Gaunt Tom R, Ben-Shlomo Yoav, Freitas Deise F, Oliveira Isabel O, Gentry-Maharaj Aleksandra, Fourkala Evangelia, Lawlor Debbie A, Hingorani Aroon D

**Author notes:** Joint last authors (Lawlor Debbie A, Hingorani Aroon D). Corresponding author: Borges MC. Rua Marechal Deodoro, 1160 - 3° Piso, Centro, Pelotas, RS, Brazil. Zipcode: 96020-220.

## Abstract

**Background:** Adiponectin, a circulating adipocyte-derived protein has insulin-sensitizing, anti-inflammatory, anti-atherogenic, and cardiomyocyte-protective properties in animal models. However, the systemic effects of adiponectin in humans are unknown.

**Objectives:** Our aims were to define the metabolic profile associated with higher blood adiponectin concentration and investigate whether variation in adiponectin concentration affects the systemic metabolic profile.

**Methods:** We applied multivariable regression in up to 5,906 adults and Mendelian randomization (using cis-acting genetic variants in the vicinity of the adiponectin gene as instrumental variables) for analysing the causal effect of adiponectin in the metabolic profile of up to 38,058 adults. Participants were largely European from six longitudinal studies and one genome-wide association consortium.

**Results:** In the multivariable regression analyses, higher circulating adiponectin was associated with higher HDL lipids and lower VLDL lipids, glucose levels, branched-chain amino acids, and inflammatory markers. However, these findings were not supported by Mendelian randomization analyses for most metabolites. Findings were consistent between sexes and after excluding high risk groups (defined by age and occurrence of previous cardiovascular event) and one study with admixed population.

**Conclusion:** Our findings indicate that blood adiponectin concentration is more likely to be an epiphenomenon in the context of metabolic disease than a key determinant.

## INTRODUCTION

The recognition that adipose tissue is an endocrine organ raised new prospects for discovering adipose-derived products that could be valuable drug targets for the treatment and prevention of cardio-metabolic diseases. In this context, adiponectin, a 30KDa protein largely produced by mature adipocytes, has been attracting widespread attention due to insulin-sensitizing, anti-inflammatory, anti-atherogenic, and cardiomyocyte-protective properties demonstrated in animal models (1).

However, human studies have yielded a far more complicated picture. Unlike most other adipokines, circulating adiponectin concentration is higher with lower adiposity (2). In prospective observational studies in humans using multivariable regression, higher circulating adiponectin is associated with lower risk of type 2 diabetes (3), hepatic dysfunction (4), and metabolic syndrome (5), but higher mortality in patients with kidney disease, heart failure, previous cardiovascular disease or general elderly cohorts (6-9); this different direction of effect between risk of incident disease and mortality among high risk groups has been called “the adiponectin paradox” (10).

Given the complex metabolic derangements that might participate in and compensatory changes that might occur in response to human diseases, the association between adiponectin concentration and cardio-metabolic biomarkers and disease end-points might be explained by reverse causality (where disease status could alter adiponectin concentration) or residual confounding (where adiponectin could be a marker of another causal factor, such as adiposity or insulin resistance) (11). Classical multivariable regression studies cannot distinguish causal from non-causal associations, and randomized controlled trials (RCTs) specifically targeting adiponectin are not possible in the absence of a specific therapeutic targeting adiponectin concentration or function.

Mendelian randomization uses genetic variants (mostly single nucleotide polymorphisms (SNPs)) that are robustly related to the risk factor of interest as tools to assess its role in causing disease (12). The random allocation of parental alleles at meiosis should theoretically reduce confounding in genetic association studies and this has been shown to be the case (13); the unidirectional flow of biological information from genetic variant to phenotypes avoids reverse causality. Mendelian randomization has been used in clinical research to investigate potential etiological mechanisms, such as the causal effects of low density lipoprotein cholesterol (LDL-c) (14), systolic blood pressure (SBP) (14) and C reactive protein (15) on coronary heart disease (CHD), validate and prioritize novel drug targets, such as interleukin-6 receptor (16), and increase understanding of current therapies, for example statins (17).

Previous Mendelian randomization studies indicate that circulating adiponectin is a consequence of low insulin sensitivity (18), but whether adiponectin concentration is also a cause of insulin sensitivity is uncertain (18-20). Using Mendelian randomization in a study of 63,746 CHD cases and 130,681 controls we have recently shown that adiponectin may not be causally related to CHD (21). Whilst multivariable analyses show higher adiponectin concentration is associated with lower glycated haemoglobin, insulin, triglycerides and higher high density lipoprotein-cholesterol (HDL-c), using Mendelian randomization, we found little evidence these were causal (21). Whether adiponectin is associated with systemic metabolic profile, and, if it is, what aspects of these associations are causal is unknown. A broader interrogation of the metabolic effects of adiponectin through high-throughput profiling of metabolic status could provide valuable insights into whether adiponectin is a non-causal biomarker or causally important in the pathophysiology of some human diseases (22).

We combined genotype, adiponectin and metabolomics profile data from six longitudinal studies and one genome-wide association consortium with the aim of (i) defining the metabolic effects of blood adiponectin concentration and (ii) investigating whether variation in adiponectin concentration is causally related to the systemic metabolic profile.

## METHODS

### Study Populations

The metabolic profile associated with blood adiponectin concentration was examined from seven data sources: the 1982 Pelotas Birth Cohort (PEL82), including adults aged 30 years old born in the city of Pelotas, Brazil, in 1982 (23, 24); the British Women’s Heart and Health Study (BWHHS), including UK women aged 60-79 years old at recruitment in 2000 (25); the Whitehall II Study (WHII), including UK government workers aged 45-69 years at phase 5 clinical assessment in 1997-1999 (26); the Caerphilly Prospective Study (CaPS), including men aged 52-72 years at phase III in 1989-1993 (27); a case-control study nested in The United Kingdom Collaborative Trial of Ovarian Cancer Screening (UKCTOCS), including UK postmenopausal women aged 50-74 years at recruitment in 2001-2005 (28); the cohort of mothers from The Avon Longitudinal Study of Children and Parents (ALSPAC-M), including UK women aged 34-63 years old at clinical assessment in 2009-2011 (29); and a metabolomics genome-wide association consortium (Metabolomics consortium), including European adults with mean age of 45 years old from 14 cohorts (30). Individual level data was available to investigators from PEL82, BWHHS, WHII, CaPS, UKCTOCS and ALSPAC-M and summary level data is publicly available from the Metabolomics consortium (URL: http://www.computationalmedicine.fi/data/NMR_GWAS/). All study participants provided written informed consent, and study protocols were approved by the local ethics committees (ethical approval for ALSPAC was also obtained from the ALSPAC Ethics and Law Committee). Studies’ characteristics are summarized on **Table 1**. We examined (possibly causal) associations of adiponectin with systemic metabolic profiles using two approaches – conventional multivariable regression and Mendelian randomization analyses. Studies must have both adiponectin and measures of some of the outcomes (but do not need genetic data) to contribute to multivariable regression analyses, and must have relevant genetic variants and outcomes (but do not need adiponectin concentration data) to contribute to Mendelian randomization analyses. **Figure 1** shows how the different data sources contributed to the two approaches.

**Figure 1.**
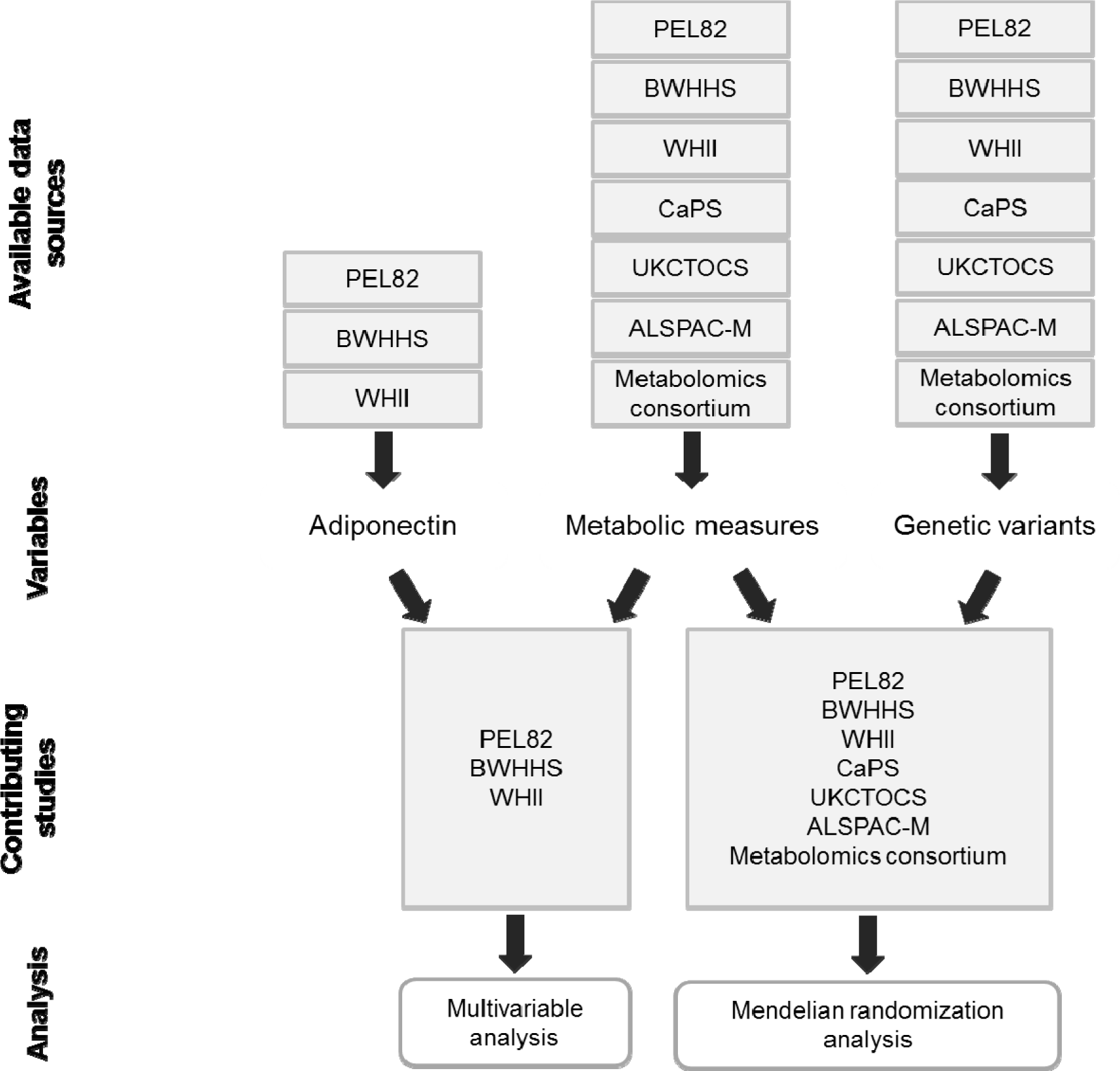
Schematic representation of studies contributing to each analytical approach. From the available data sources, three had data on adiponectin and metabolic measures and could contribute to multivariable analysis (PEL82, BWHHS and WHII) and all had data on genetic variants and metabolic measures and could contribute to Mendelian randomization analysis (PEL82, BWHHS, WHII, CaPS, UKCTOCS, ALSPAC-M, and Metabolomics consortium). ALSPAC-M: The Avon Longitudinal Study of Children and Parents – mothers’ cohort; BWHHS: British Women’s Heart and Health Study; CaPS: The Caerphilly Prospective Study; PEL82: 1982 Pelotas Birth Cohort; UKCTOCS: case-control study nested in The United Kingdom Collaborative Trial of Ovarian Cancer Screening; WHII: Whitehall-II Study.

**Table 1.**
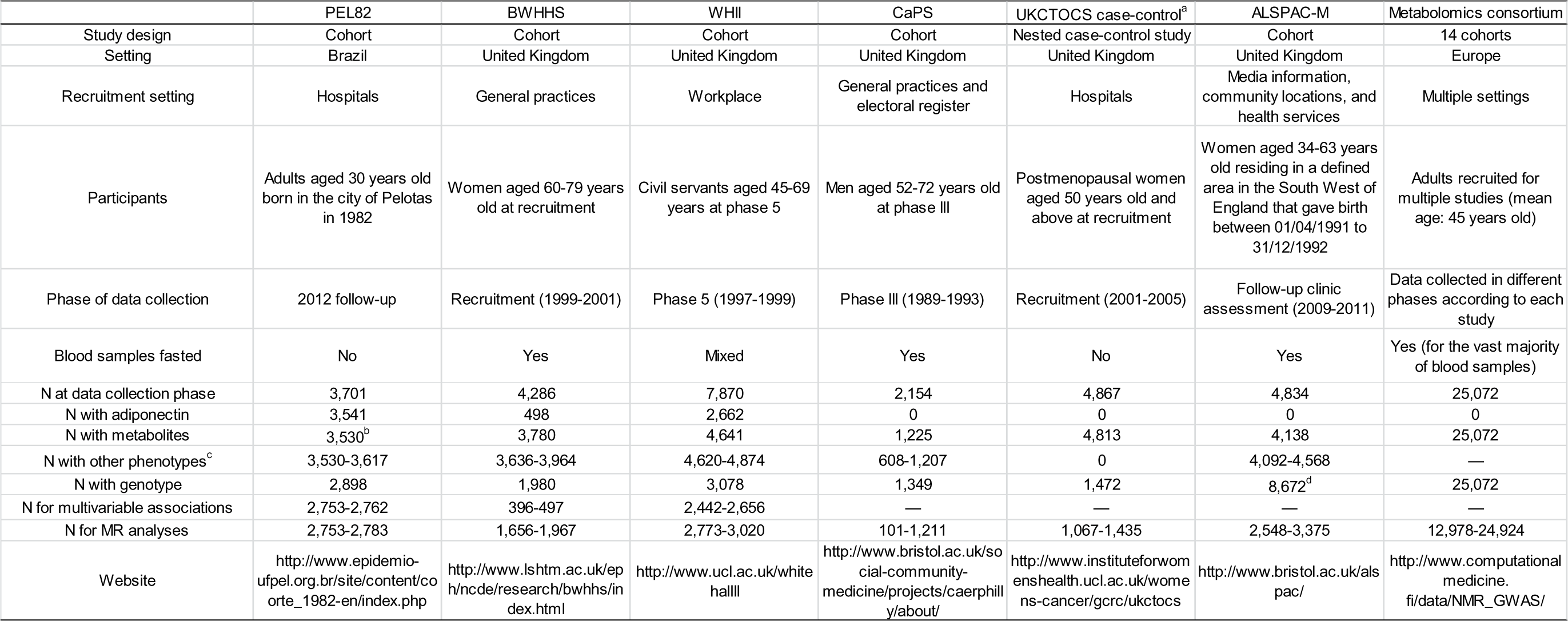
Characteristics of participating studies.

### Metabolite Quantification

A high-throughput serum nuclear magnetic resonance (NMR) spectroscopy platform was utilized to quantify up to 150 metabolic measures and 83 derived measures (ratios) in each study. This NMR platform has been used in several other studies (22, 31, 32) and methodological details have been described elsewhere (33, 34). 66 out of 150 metabolic measures were selected for this study aimed at broadly representing the systemic metabolite profile, as previously reported by Wurtz et al (35), including: lipoprotein traits (lipid content, particle size, and apolipoproteins), free fatty acids, amino acids, glycolysis-related metabolites, ketone bodies, fluid balance (albumin and creatinine), and inflammatory markers (glycoprotein acetyls). The remaining 84 metabolic measures from the NMR platform are related to other lipid fractions (esterified and free cholesterol, total cholesterol, triglycerides, and phospholipids) and particle concentration from 14 lipoprotein subclasses and are not presented in this study. Instead, we present the total lipid content of each of the 14 lipoprotein subclasses, which is highly correlated to their respective lipid fractions and particle concentration and comprehensively represents the plasma lipid partitioning across lipoproteins. Eight additional measures, not obtained from the NMR platform, were also included: C-reactive protein (CRP), interleukin (IL)-6, fibrinogen, blood viscosity, insulin, glycated haemoglobin (Hb_A1c_), and systolic (SBP) and diastolic blood pressure (DBP). PEL82 did not have data on metabolic measures from NMR platform and contributed data to analyses of conventional lipid risk factors (total cholesterol, HDL-c, LDL-c, and triglycerides (TG)), and some of the additional measures described (CRP, Hb_A1c_, SBP, DBP). Adiponectin was assayed using an enzyme-linked immunosorbent assay (ELISA) in PEL82, BWHHS and WHII. Data on adiponectin level was not available from CaPS, UKCTOCS, ALSPAC-M, and the Metabolomics consortium. Blood samples used for adiponectin, NMR metabolites, and other blood based outcomes were taken after overnight or minimum 6-hours fast in BWHHS, CaPS, and ALSPAC-M and on non-fasting samples in PEL82 and UKCTOCS. In WHII, participants attending the morning clinic were asked to fast overnight and those attending in the afternoon were asked to have a light, fat-free breakfast before 0800 hours. The vast majority of samples contributing to the Metabolomics consortium were fasting samples.

### Genotyping

BWHHS, CaPS, WHII and UKCTOCS participants were genotyped using Metabochip, a platform comprising 200,000 SNPs, which cover the loci identified by GWAS in cardio-metabolic diseases, and rare variants from the 1000 Genomes Project (36). Quality control criteria and imputation using 1000 Genomes European ancestry reference samples have been previously described for studies within UCLEB consortium (37). In ALSPAC-M, 557,124 SNPs were directly genotyped using Illumina human660W quad. For quality control, SNPs were excluded if missingness > 5%, Hardy-Weinberg equilibrium P-value < 1*10^−6^ or minor allele frequency < 1%, and samples were excluded if missingness > 5%, indeterminate X chromosome heterozygosity, extreme autosomal heterozygosity or showing evidence of population stratification. Imputation was performed using 1000 genomes reference panel (Phase 1, Version 3) (phased using ShapeIt v2. r644, haplotype release date Dec 2013) and Impute V2.2.2. For PEL82, genotyping was performed by using the Illumina HumanOmni2.5-8v1 array (Illumina Inc.) and approximately 2,500,000 SNPs were genotyped (38). For PEL82, quality control criteria have been previously described (38) and imputation was performed in two steps: first, genotypes were phased using SHAPEIT; then, IMPUTE2 was used for the actual imputation. For autosomal and X-chromosome SNPs, 1000 Genomes Phase I integrated haplotypes (December 2013 release) and 1000 Genomes Phase I integrated variant set (March 2012 release), respectively, were used. For PEL82, ancestry-informative principal components were based on 370,539 SNPs shared by samples from the HapMap Project, the Human Genome Diversity Project (HGDP), and PEL82. The following HapMap samples were used as external panels: 266 Africans, 262 Europeans (American and Italian), 77 admixed Mexican Americans, 83 African Americans, and 93 Native Americans from the HGDP [more details can be found in (39)]. Cohorts contributing to the Metabolomics consortium used different SNP arrays, non-genotyped SNPs were imputed using a 1000 Genomes Project March 2012 version and SNPs with accurate imputation (proper info > 0.4) and minor allele count >3 were combined in fixed-effects meta-analysis using double genomic control correction. Further details can be found in the consortium publication (30).

### Other covariates

Anthropometric variables (weight and height) were measured in each study using standard procedures and body mass index (BMI) was calculated as weight (kg)/height (m)^2^. Demographic and smoking status information were obtained through questionnaires.

### Data analysis

Prior to analyses, metabolic measures were adjusted for age, sex, and, if applicable, place of recruitment (BWHHS and UKCTOCS) or principal components of genomic ancestry (PEL82 and some studies contributing to Metabolomics consortium) and the resulting residuals were transformed to normal distribution by inverse rank-based normal transformation. Pregnant women from PEL82 (n = 73) and ALSPAC-M (n = 12) were excluded. As the 74 analysed metabolites are highly correlated, we adopted a similar strategy to the Metabolomics consortium (30) to correct for multiple testing by estimating the number of independent tests as the number of principal components that explained over 95% of variance in metabolites concentration using data from the two studies (BWHHS and WHII) with the largest available number of metabolites (n = 27 principal components in both studies). As a result, for both multivariable and Mendelian randomization analyses, we corrected for multiple testing using the Bonferroni method considering 27 independent tests (p = 0.05 ÷ 27 ≈ 0.0019).

#### Multivariable regression analysis

The conventional multivariable regression association of adiponectin with individual metabolites was estimated using a two-stage individual participant meta-analysis. In the first stage, linear regression models were fitted for each study. In the second stage, study-specific estimates were meta-analysed using DerSimonian & Laird random effect model (40). Heterogeneity across studies was assessed using I^2^ (as a measure of the relative size of between-study variation and within-study error) (41). Three types of subgroup analyses were conducted: sex-stratified analysis, analysis excluding individuals with high risk of cardiometabolic disease (those that had experienced coronary artery disease or stroke or those older than 65 years old) and analysis restricted to European studies (excluding PEL82).

#### Genetic analyses

##### Selection of genetic variants

The SNPs used for the Mendelian randomization analysis were selected from 145 SNPs with good evidence (p < 5*10^−8^) for association with blood adiponectin concentration in the European ancestry GWAS meta-analysis from the ADIPOGen consortium (42). Independent SNPs within the *ADIPOQ* locus (± 50 kb) have been previously selected by Dastani et al (2013) (43) by linkage disequilibrium (LD) prunning of the genome-wide significant SNPs, retaining SNPs that explained most variance in adiponectin concentration in each LD block (LD threshold: R^2^ < 0. 05 in HapMap CEU population (Utah residents with Northern and Western European ancestry)). This resulted in four SNPs (rs6810075, rs16861209, rs17366568, and rs3774261), which are estimated to explain approximately 4% of variance in adiponectin concentration (**Table 2 and Supplementary methods**). Data for the association of each selected SNP with adiponectin concentration in the discovery GWAS sample was downloaded from ADIPOGen website (https://www.mcgill.ca/genepi/adipogen-consortium).

**Table 2.** Characteristics of studies’ populations.

|  | PEL82 | BWHHS | WHII | CaPS | UKCTOCS | ALSPAC-M | Metabolomics consortium |
| --- | --- | --- | --- | --- | --- | --- | --- |
|  | % |  |  |  |  |  |  |
| Male | 49 | 0 | 72 | 100 | 0 | 0 | 45 |
| White | 75 | 100 | 93 | 100 | 97 | 97 | NA<sup>a</sup> |
| Smoker | 24 | 12 | 17 | 20 | — | 11 | NA |
| Overweight/obese | 58 | 72 | 57 | 69 | 60 | 56 | NA |
|  | Median (p25, p75) |  |  |  |  |  |  |
| Age (years) | 30 (30, 30) | 69 (64, 73) | 55 (51, 61) | 56 (53, 60) | 66 (60, 70) | 48 (45, 51) | 45 (24, 61)<sup>b</sup> |
| Adiponectin (µg/mL) | 7.9 (5.2, 11.9) | 15.8 (10.8, 21.5) | 8.5 (6.1, 12) | — | — | — | — |
| Glucose (mmol/L) | 4.8 (4.4, 5.3) | 4.7 (4.3, 5.1) | 5 (4.7, 5.4) | 3.8 (3.5, 4.2) | 2.2 (1.7, 3.1) | 4.4 (4.1, 4.7) | NA |
| HDL-c (mmol/L) | 1.5 (1.2, 1.7) | 1.6 (1.4, 1.9) | 1.5 (1.3, 1.7) | 0.9 (0.7, 1) | 1.6 (1.4, 1.9) | 1.7 (1.5, 1.9) | NA |
| LDL-c (mmol/L) | 2.7 (2.3, 3.3) | 2.3 (1.9, 2.8) | 1.9 (1.6, 2.2) | 1.6 (1.3, 1.9) | 1.8 (1.4, 2.2) | 1.5 (1.2, 1.8) | NA |
| TG (mmol/L) | 1.1 (0.8, 1.6) | 1.5 (1.1, 2) | 1.1 (0.9, 1.5) | 1.5 (1.2, 2) | 1.5 (1.1, 2.1) | 0.9 (0.7, 1.2) | — |
| SBP (mmHg) | 120 (112, 130) | 146 (130, 163) | 121 (111, 133) | 144 (130, 160) | — | 117 (110, 125) | — |
| DBP (mmHg) | 75 (69, 81) | 79 (71, 87) | 77 (70, 84) | 84 (76, 92) | — | 71 (66, 77) | — |

<sup>a</sup> Cohorts contributing to the Metabolomics consortium were of European origin
<sup>b</sup> Overall mean age (and range of mean age across studies)
ALSPAC-M: The Avon Longitudinal Study of Children and Parents – mothers' cohort; BWHHS: British Women's Heart and Health Study; CaPS: The Caerphilly Prospective Study; DBP: diastolic blood pressure; HDL-c: high-density lipoprotein-cholesterol; LDL-c: low-density lipoprotein-cholesterol; PEL82: 1982 Pelotas Birth Cohort; SBP: systolic blood pressure; TG: triglycerides; UKCTOCS: case-control study nested in The United Kingdom Collaborative Trial of Ovarian Cancer Screening; WHII: Whitehall-II Study.

##### Association of genetic variants with classical confounders

The association between genetic variants and classical confounders [sex, age, ancestry (European vs non European), current smoking (yes vs no), and body mass index] was examined for each study using logistic or linear regression models for binary or continuous variables, respectively.

##### Mendelian randomization analysis

In order to allow all participants with relevant genetic and metabolic measure data to contribute to analyses, even when adiponectin data was not available (as in CaPS, UKCTOCS, ALSPAC-M, and Metabolomics consortium), a two-sample Mendelian randomization design was used, in which data for the association between genetic variants and adiponectin levels were obtained from an external data source, the ADIPOGen consortium (42). The two-sample Mendelian randomization is a recent extension to the more conventional one-sample Mendelian randomization and has the additional advantage of avoiding bias due to genetic variants correlating with confounders by chance (statistical overfitting) when samples are independent (44). The two-sample Mendelian randomization estimates and respective standard errors were obtained using the inverse variance-weighted (IVW) method, as described by Burgess et al. (45) and detailed in Supplementary Methods. Study-specific Mendelian randomization estimates were meta-analysed using DerSimonian & Laird random effect model (40). Heterogeneity across studies was assessed using I^2^ (41). Subgroup analyses were conducted considering individual-level (sex and risk of cardiometabolic disease) and study-level characteristics (European vs non-European studies). The Metabolomics consortium did not contribute to subgroup analysis of individual-level characteristics as only summary data was available. Results from conventional multivariable and Mendelian randomization analyses were compared by using the Z-test for each metabolic measure (details in the Supplementary methods) and by estimating the correlation between multivariable and Mendelian randomization estimates across all metabolic measures. Power calculations for Mendelian randomization analysis are available in **Supplementary table 1**.

## RESULTS

The study included a median sample size of 3,006 adults in the multivariable analysis (range: 2,497-5,906) and a median sample size of 23,884 adults in the Mendelian randomization analysis (range: 4,645-38,058). Characteristics of participants from each contributing study are listed in **Table 2**.

### Adiponectin and the Systemic Metabolic Profile

In the multivariable analysis, adiponectin was associated with 59 out of 74 (80%) metabolites at nominal level (p < 0.05) and 49 out of 74 (66%) after correcting for multiple testing (p < 0.0019). Overall, higher circulating adiponectin was associated with a healthier systemic metabolite profile. Blood adiponectin concentration was strongly related to multiple lipoprotein traits. With higher adiponectin concentration, lipid concentration was lower in VLDL subclasses and higher in HDL subclasses, except for small HDL. There was no strong evidence of circulating adiponectin associating with total lipid content in LDL subclasses or in IDL, although adiponectin concentration was inversely associated with LDL-cholesterol. Higher adiponectin was associated with lower concentration of cholesterol and triglycerides, lower mean particle diameter in VLDL and higher cholesterol concentration and mean particle diameter in HDL. Higher adiponectin concentration was also associated with higher concentration of apolipoprotein (Apo)-AI and phospholipids and lower concentration of triglycerides and diglycerides (**Figure 2**).

**Figure 2.**
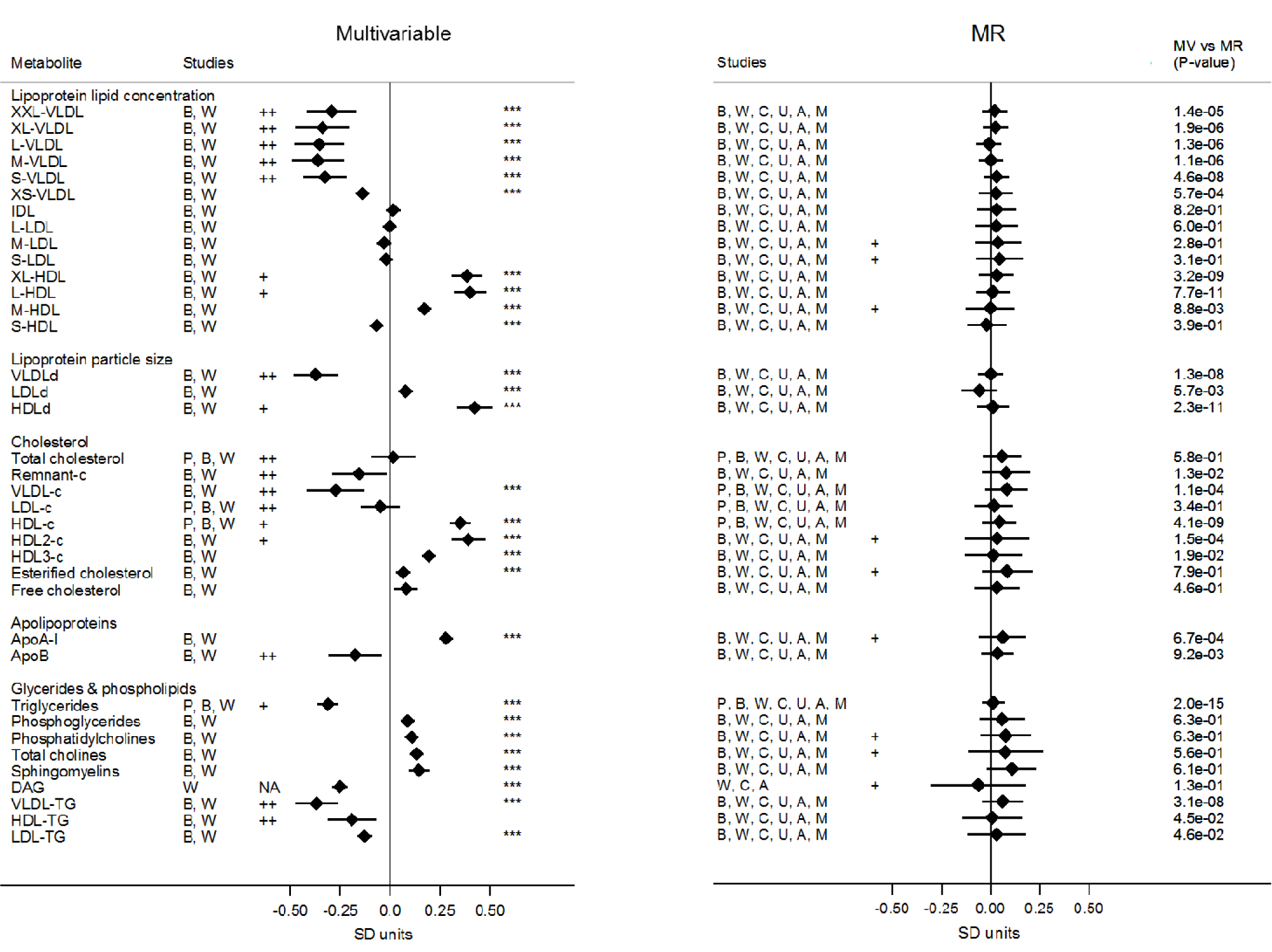
Association of lipoprotein traits with blood adiponectin levels from observational and Mendelian randomization (MR) analysis. Values are expressed as units of standardized log metabolite concentration (and 95% CI) per 1 unit increment of standardized log adiponectin levels. P-values for the association between adiponectin and metabolites are indicated by three asterisks (“***”) if lower than Bonferroni-adjusted threshold (P-value < 0.00068). Heterogeneity was considered substantial if I^2^ = 50-75% (“+”) or very high if I^2^ > 75% (“++”). P-values for the comparison between multivariable and Mendelian randomization estimates are displayed in the column “MR vs MV (P-value)”. Metabolic measures were adjusted for age, sex, and, if applicable, place of recruitment (BWHHS and UKCTOCS) or principal components of genomic ancestry (PEL82 and some studies contributing to Metabolomics consortium) and the resulting residuals were transformed to normal distribution by inverse rank-based normal transformation. XXL: extremely large, XL: very large, L: large, M: medium, S: small, XS: very small, VLDL: very low-density lipoprotein, LDL: low-density lipoprotein, IDL: intermediate-density lipoprotein, HDL: high-density lipoprotein, c: cholesterol, DAG: diglycerides, TG: triglycerides, P: 1982 Pelotas Birth Cohort, B: British Women Heart and Health Study, W: Whitehall II Study, C: The Caerphilly Prospective Study, U: UKCTOCS nested case-control study, A: The Avon Longitudinal Study of Children and Parents – mothers’ cohort, M: Metabolomics consortium, SD units: standard deviation units, CI: confidence interval.

Higher circulating adiponectin was also associated with healthier glycemic status (lower glucose and insulin concentration), lower blood concentration of glycolysis-related metabolites (lactate and pyruvate), saturated fatty acids, systemic inflammatory markers (C-reactive protein, fibrinogen, interleukin-6, glycoprotein acetyls and blood viscosity), systolic blood pressure, creatinine, and higher ketone bodies (acetoacetate). In addition, higher adiponectin concentration was associated with lower concentrations of free branched chain amino acids (isoleucine, leucine, and valine), aromatic amino acids (phenylalanine and tyrosine), and alanine and higher concentration of glutamine (**Figure 3**).

**Figure 3.**
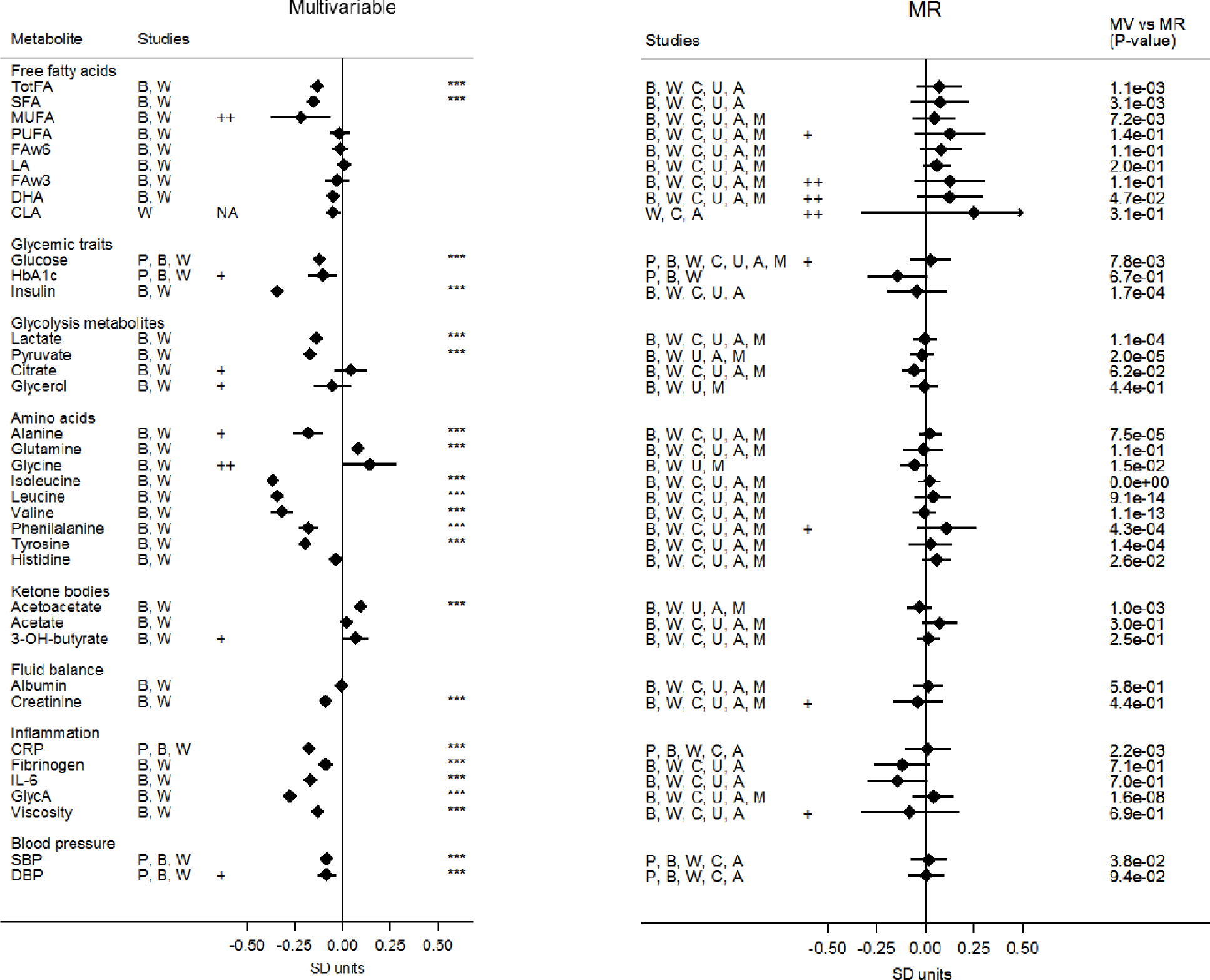
Association of multiple metabolic measures with blood adiponectin levels from observational and Mendelian randomization analysis. Values are expressed as units of standardized log metabolite concentration (and 95% CI) per 1 unit increment of standardized log adiponectin levels. P-values for the association between adiponectin and metabolites are indicated by three asterisks (“***”) if lower than Bonferroni-adjusted threshold (P-value < 0.00068). Heterogeneity was considered substantial if I^2^ = 50-75% (“+”) or very high if I^2^ > 75% (“++”). P-values for the comparison between multivariable and Mendelian randomization estimates are displayed in the column “MR vs MV (P-value)”. Metabolic measures were adjusted for age, sex, and, if applicable, place of recruitment (BWHHS and UKCTOCS) or principal components of genomic ancestry (PEL82 and some studies contributing to Metabolomics consortium) and the resulting residuals were transformed to normal distribution by inverse rank-based normal transformation. TotFA: total fatty acids, SFA: saturated fatty acid, MUFA: monounsaturated fatty acid, PUFA: polyunsaturated fatty acids, FAw6: omega-6 fatty acid, LA: linoleic acid, FAw3: omega-3 fatty acid, DHA: docosaexaenoic acid, CLA: conjugated linoleic acids, HbA1c: glycated haemoglobin, CRP: c-reactive protein, IL-6: interleukin-6, GlycA: glycoprotein acetyls, SBP: systolic blood pressure, DBP: diastolic blood pressure, P: 1982 Pelotas Birth Cohort, B: British Women Heart and Health Study, W: Whitehall II Study, C: The Caerphilly Prospective Study, U: UKCTOCS nested case-control study, A: The Avon Longitudinal Study of Children and Parents – mothers’ cohort, M: Metabolomics consortium, SD units: standard deviation units, CI: confidence interval.

In the multivariable analyses, evidence of heterogeneity in pooled estimates across studies was substantial (I^2^ = 50%-75%) for 12 and very high (I^2^ > 75%) for 15 metabolic measures (**Figure 2** and **3** and **Supplementary table 2**). This did not seem to be accounted by sex (**Supplementary figures 1 to 4**), geographic location (**Supplementary figures 5 and 6**), or high risk of disease (**Supplementary figures 7 and 8**).

### Causal effects of adiponectin on the Systemic Metabolic Profile

Characteristics of the four SNPs (rs6810075, rs16861209, rs17366568 and rs3774261) used in Mendelian randomization and their association with adiponectin concentration are shown in **Table 3**. Overall, SNPs effect allele frequency was similar across studies. Two SNPs had lower allele frequency in the Metabolomics consortium (rs6810075: 51% vs. 65-69% in other studies; rs16861209: 5% vs. 9-11% in other studies) and one SNP had a higher frequency in PEL82 compared to other studies (rs3774261: 49% vs. 38-39% in other studies) (**Table 3**). As expected, the selected SNPs were not associated with classical confounders overall (**Supplementary table 3**).

**Table 3.**
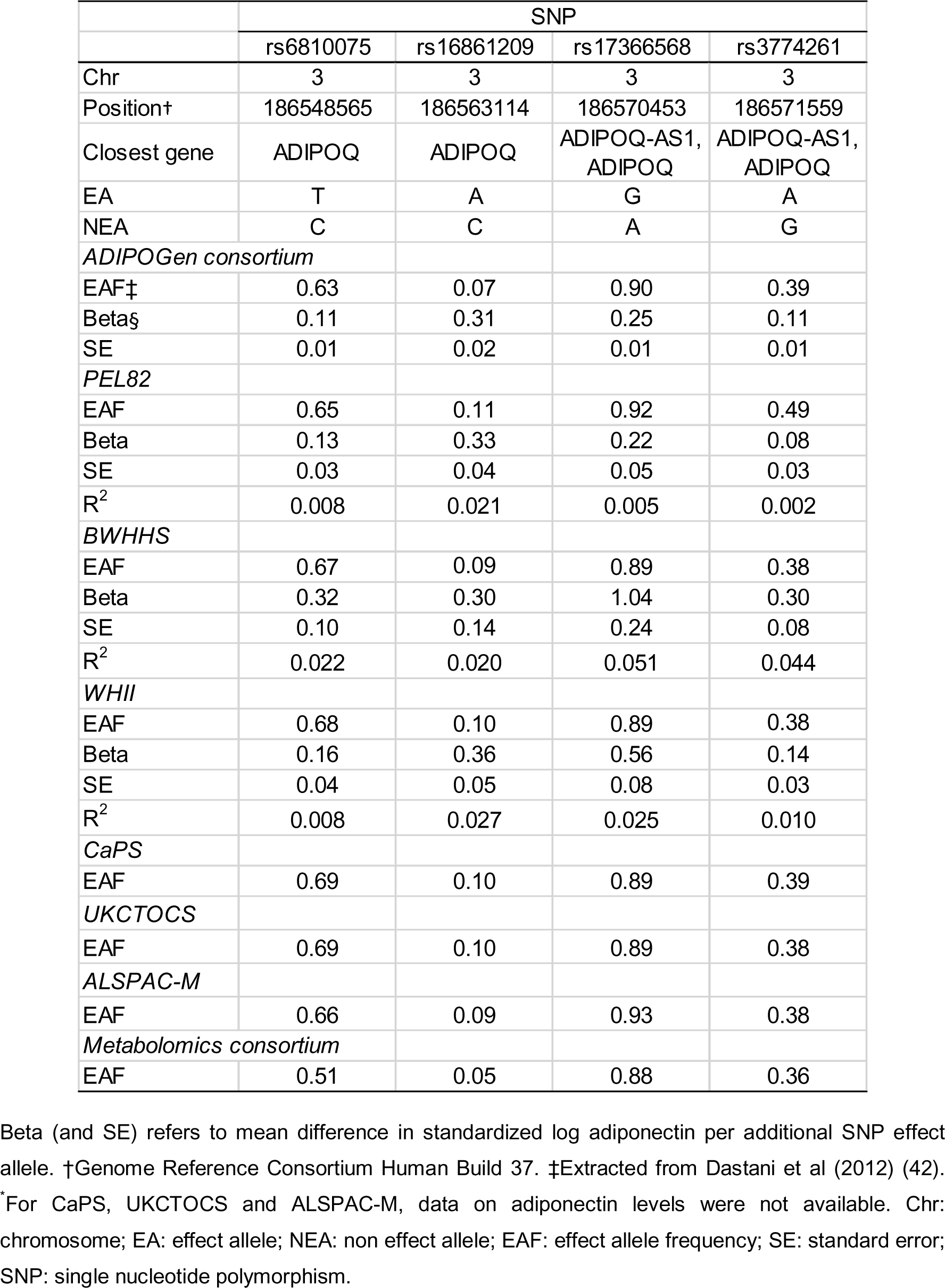
Characteristics of SNPs selected for Mendelian randomization analysis.

|  | SNP |  |  |  |
| --- | --- | --- | --- | --- |
|  | rs6810075 | rs16861209 | rs17366568 | rs3774261 |
| Chr | 3 | 3 | 3 | 3 |
| Position† | 186548565 | 186563114 | 186570453 | 186571559 |
| Closest gene | ADIPOQ | ADIPOQ | ADIPOQ-AS1,<br>ADIPOQ | ADIPOQ-AS1,<br>ADIPOQ |
| EA | T | A | G | A |
| NEA | C | C | A | G |
| *ADIPOGen consortium* |  |  |  |  |
| EAF‡ | 0.63 | 0.07 | 0.90 | 0.39 |
| Beta§ | 0.11 | 0.31 | 0.25 | 0.11 |
| SE | 0.01 | 0.02 | 0.01 | 0.01 |
| *PEL82* |  |  |  |  |
| EAF | 0.65 | 0.11 | 0.92 | 0.49 |
| Beta | 0.13 | 0.33 | 0.22 | 0.08 |
| SE | 0.03 | 0.04 | 0.05 | 0.03 |
| R<sup>2</sup> | 0.008 | 0.021 | 0.005 | 0.002 |
| *BWHHS* |  |  |  |  |
| EAF | 0.67 | 0.09 | 0.89 | 0.38 |
| Beta | 0.32 | 0.30 | 1.04 | 0.30 |
| SE | 0.10 | 0.14 | 0.24 | 0.08 |
| R<sup>2</sup> | 0.022 | 0.020 | 0.051 | 0.044 |
| *WHII* |  |  |  |  |
| EAF | 0.68 | 0.10 | 0.89 | 0.38 |
| Beta | 0.16 | 0.36 | 0.56 | 0.14 |
| SE | 0.04 | 0.05 | 0.08 | 0.03 |
| R<sup>2</sup> | 0.008 | 0.027 | 0.025 | 0.010 |
| *CaPS* |  |  |  |  |
| EAF | 0.69 | 0.10 | 0.89 | 0.39 |
| *UKCTOCS* |  |  |  |  |
| EAF | 0.69 | 0.10 | 0.89 | 0.38 |
| *ALSPAC-M* |  |  |  |  |
| EAF | 0.66 | 0.09 | 0.93 | 0.38 |
| *Metabolomics consortium* |  |  |  |  |
| EAF | 0.51 | 0.05 | 0.88 | 0.36 |

Beta (and SE) refers to mean difference in standardized log adiponectin per additional SNP effect allele. †Genome Reference Consortium Human Build 37. ‡Extracted from Dastani et al (2012) (42).
\*For CaPS, UKCTOCS and ALSPAC-M, data on adiponectin levels were not available. Chr: chromosome; EA: effect allele; NEA: non effect allele; EAF: effect allele frequency; SE: standard error; SNP: single nucleotide polymorphism.

Findings from Mendelian randomization analysis were largely inconsistent with results from multivariable analysis. Firstly, there was no evidence that adiponectin influenced HDL and VLDL traits (**Figure 2**). Secondly, genetically-increased adiponectin levels were not associated with glycemic traits, free amino acids, and glycolysis-related metabolites (**Figure 3**). Results were less conclusive for some inflammatory markers (IL-6 and fibrinogen) (**Figure 3**). Thirdly, there was strong statistical evidence that associations from multivariable and Mendelian randomization analyses were inconsistent with each other (**Figure 2** and **Figure 3**) and the overall correlation between multivariable and Mendelian randomization estimates was very low (r = 0.10) (**Figure 4**). Finally, in the Mendelian randomization analysis, adiponectin was not associated with any of the metabolic analyses at either p < 0.05 or p < 0.00068.

**Figure 4.**
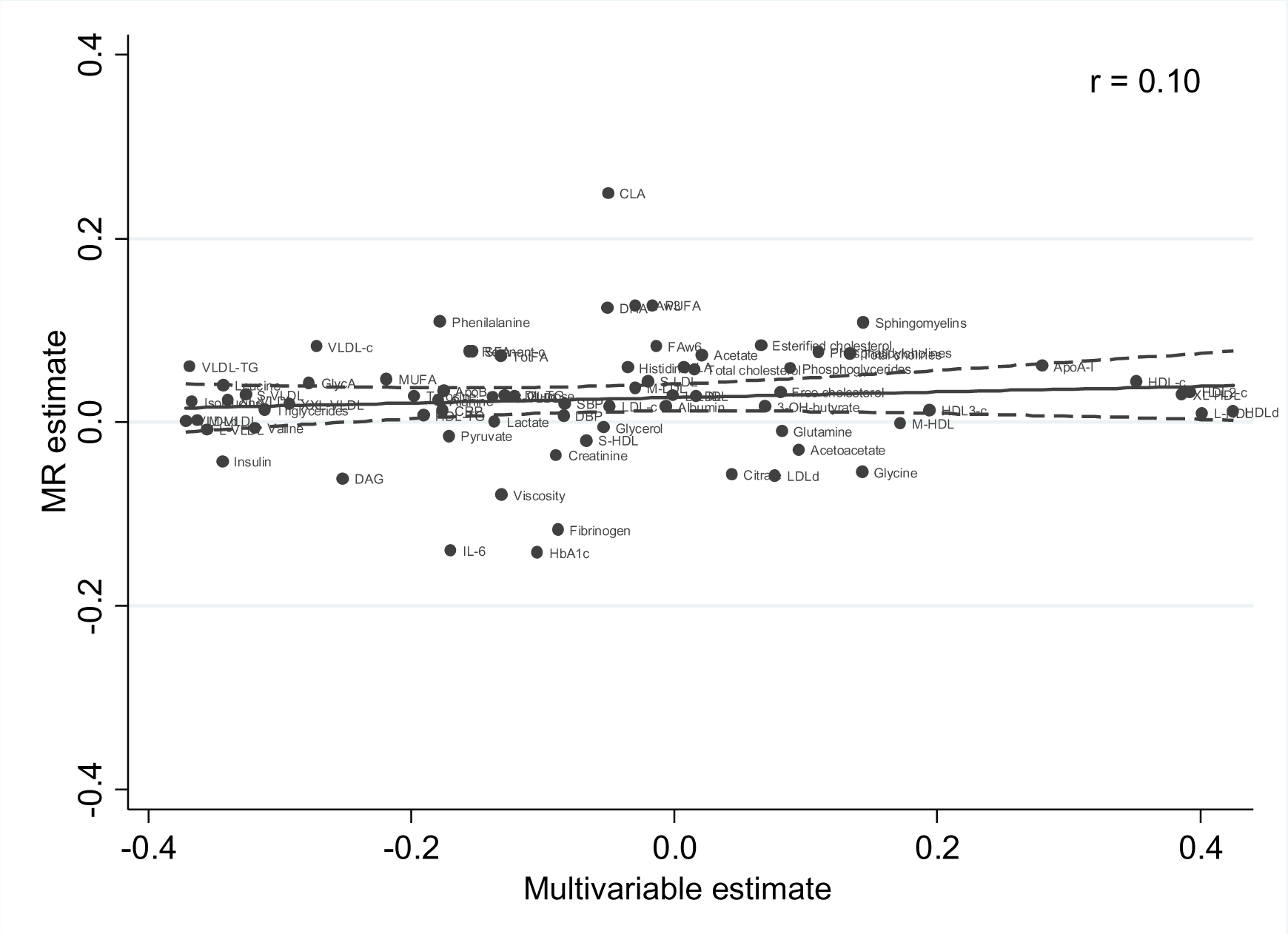
Correlation between estimates from multivariable regression and Mendelian randomization (MR). XXL: extremely large, XL: very large, L: large, M: medium, S: small, XS: very small, VLDL: very low-density lipoprotein, LDL: low-density lipoprotein, IDL: intermediate-density lipoprotein, HDL: high-density lipoprotein, c: cholesterol, DAG: diglycerides, TG: triglycerides, SFA: saturated fatty acid, MUFA: monounsaturated fatty acid, FAw: 6: omega-6 fatty acid, LA: linoleic acid, DHA: docosaexaenoic acid, FAw3: omega-3 fatty acid, HbA1c: glycated haemoglobin, CRP: c-reactive protein, IL-6: interleukin-6, GlycA: glycoprotein acetyls, SBP: systolic blood pressure, DBP: diastolic blood pressure, r: Pearson correlation coefficient.

In the Mendelian randomization analyses, evidence of heterogeneity in pooled estimates across studies were substantial (I^2^ = 50%-75%) for 14 and very high (I^2^ > 75%) for 3 metabolic measures, suggesting lower heterogeneity in models from genetic analysis than from the multivariable analyses (**Figure 2** and **3** and **Supplementary table 2**). This did not seem to be driven by sex differences (**Supplementary figures 1 to 4**), geographic location/ethnicity (**Supplementary figures 5 and 6**), or high risk of disease (**Supplementary figures 7 and 8**).

## DISCUSSION

In up to 5,906 adults we found using multivariable regression analyses that circulating adiponectin was associated with a pattern of systemic metabolites levels associated with good health. Higher blood adiponectin concentration was associated with higher HDL lipids and lower VLDL lipids, glycaemia, and branched-chain amino acids levels. However, when we used genetic variants in the *ADIPOQ* locus to test the causal effect of adiponectin on systemic metabolic profiles amongst up to 38,058 adults, we found little evidence that the associations were causal.

Despite the evidence of shared genetic architecture between adiponectin concentration and cardio-metabolic diseases (42), previous Mendelian randomization studies have cast doubt on the causal role of blood adiponectin levels in the risk of type 2 diabetes (18) and coronary heart disease (21). In addition, there seems to be no consistent evidence that circulating adiponectin causally affects traditional cardiovascular risk factors, such as HDL-c, LDL-c, triglycerides, and fasting glucose in the population (18). We have added importantly to those previous studies and explored effects on systemic metabolic profiles. Taken together, this and previous Mendelian randomization studies suggest that the association between circulating adiponectin and metabolic biomarkers and cardio-metabolic diseases is likely to be explained by shared factors (confounding) rather than by a direct role of adiponectin on metabolism and downstream cardio-metabolic disease. These results are in contrast to findings from animal models pointing to insulin-sensitizing, and antiatherogenic actions of adiponectin (1).

Circulating adiponectin is known to be substantially reduced among obese individuals, particularly in the presence of central fat accumulation (46). A recent Mendelian randomization study examining the causal metabolic effects of BMI demonstrated that lower BMI was related to favorable lipoprotein subclass profile and lower concentration of branched-chain amino acids, inflammatory markers, and insulin (35), which is remarkably similar to our results from the conventional multivariable analysis. In addition, numerous studies have shown that adiponectin production is supressed by insulin action in humans, which seems to be at least partly attributed to regulation at the transcriptional level (11, 47). As an example, elevated circulating adiponectin is found in contexts of both primary deficiency of insulin (type 1 diabetes) (48) and global insulin resistance due to genetic or acquired defects in the insulin receptor (49). Evidence from animal models has raised the possibility of a bidirectional relationship between adiponectin and insulin concentration (50). Early Mendelian randomization studies did indicate that adiponectin could mitigate insulin resistance (19, 20); however, these results could not be replicated in a larger Mendelian randomization study (18), as well as in our study presented here. The well-known metabolic effects of adiposity and insulin on circulating adiponectin concentration reinforce that the clustering of adiponectin and several traditional and novel biomarkers is likely to result from confounding due to increasing adiposity and disruption of insulin action.

Strengths of our study include detailed metabolic profile in several longitudinal studies, which enabled us to characterize the metabolic profile of high adiponectin concentration beyond traditional biomarkers, as well as the use of Mendelian randomization to disentangle the causal effect of adiponectin on the metabolism. Mendelian randomization analysis can reliably test for the presence of a causal relation under the three assumptions of an instrumental variable that the genetic variants are robustly associated with the risk factor of interest (adiponectin) (1), should only affect the outcome (metabolites) through the exposure (2), and are not associated with exposure-outcome confounders (3) (51). To ensure that IV assumptions were met, or were at least plausible, we only used SNPs strongly and specifically (within *ADIPOQ* gene) related to adiponectin concentration as instrumental variables and we adjusted for population structure in models using data from PEL82 to avoid confounding by population stratification. One of the limitations of our study was the limited power in subgroup analyses including only individual-level data (sex- and risk-stratified analyses), which limited our investigation of potential sources of heterogeneity. Another limitation was the absence of data on high-molecular weight adiponectin, which is believed to account for most of the adiponectin biological effects in experimental settings. However, most human (and many animal model) studies have not used high-molecular weight adiponectin, and we found the same multivariable observational associations as in previous studies.

Overall, our findings suggest that altered total blood adiponectin concentration is an epiphenomenon in the context of metabolic disease, rather than a key determinant. Therefore, interventions targeting manipulation of adiponectin concentration are unlikely to result in therapeutic benefits for tackling cardiovascular diseases. Our results highlight the potential of Mendelian randomization analysis and high-throughput metabolomics profiling to yield important insights to advance our understanding in the pathophysiology of common complex diseases and to inform which targets are ‘best-bets’ for taking forward into drug development, given that drug target validation is a key obstacle underlying the unsustainably high rate of drug development failure. Whilst our, and other studies, suggest adiponectin is not a valuable target for developing drugs aimed at preventing cardio-metabolic diseases, it may nonetheless be a valuable biomarker for predicting these diseases given the wide ranging associations shown here. The associations we have found would need to be replicated in additional independent studies before testing their ability to predict disease outcomes.

## Acknowledgements

We acknowledge Andy Ryan for his contribution to data collection from UKCTOCS. MCB, DLSF, DAL and TRG work in the MRC Integrative Epidemiology Unit at the University of Bristol that receives funding from the UK Medical Research Council (MC_UU_12013/5 and MC_UU_12013/8). DAL is a UK National Institute of Health Research Senior Investigator (NF-SI-0611-10196). MKiv is supported by the UK Medical Research Council (K013351). **PEL82** (1982 Pelotas Birth Cohort) is conducted by Postgraduate Program in Epidemiology at Universidade Federal de Pelotas with the collaboration of the Brazilian Public Health Association (ABRASCO). From 2004 to 2013, the Wellcome Trust supported PEL82. The International Development Research Center, World Health Organization, Overseas Development Administration, European Union, National Support Program for Centers of Excellence (PRONEX), the Brazilian National Research Council (CNPq), and the Brazilian Ministry of Health supported previous phases of the study. The **UCLEB** (UCL-LSHTM-Edinburgh-Bristol) consortium, which is supported by BHF Programme Grant RG/10/12/28456, consists of 12 studies: Northwick Park Heart Study II (NPHS II), British Regional Heart Study (BRHS), Whitehall II Study (WHII), English Longitudinal Study of Ageing (ELSA), Medical Research Council National Survey of Health and Development (MRC NSHD), 1958 Birth cohort (1958BC), Caerphilly prospective study (CaPS), British Women’s Heart and Health Study (BWHHS), Edinburgh Artery Study (EAS), Edinburgh Heart Disease Prevention Study (EHDPS), Edinburgh Type 2 Diabetes Study (ET2DS) and Asymptomatic Atherosclerosis Aspirin Trial (AAAT). **BWHHS** is supported by funding from the British Heart Foundation and the Department of Health Policy Research Programme (England). EAS is funded by the British Heart Foundation (Programme Grant RG/98002), with Metabochip genotyping funded by a project grant from the Chief Scientist Office of Scotland (Project Grant CZB/4/672). The **WHII study** is supported by grants from the Medical Research Council (K013351), British Heart Foundation (RG/07/008/23674), Stroke Association, the US National Heart Lung and Blood Institute (5RO1 HL036310), the US National Institute on Aging (5RO1AG13196) the US Agency for Healthcare Research and Quality (HS06516); and the John D. and Catherine T. MacArthur Foundation Research Networks on Successful Midlife Development and Socio-economic Status and Health. **CaPS** was funded by the Medical Research Council and undertaken by the former MRC Epidemiology Unit (South Wales). The CaPS DNA bank was established with funding from a MRC project grant. The CaPS data archive is maintained by the University of Bristol. MRC Integrative Epidemiology Unit, Bristol is supported by MRC grants (MR_UU_12013/1, MR_UU_12013/5 and MR_UU_12013/8). **UKCTOCS** was funded by the Medical Research Council (G9901012 and G0801228), Cancer Research UK (C1479/A2884), and the Department of Health, with additional support from The Eve Appeal. Phenotypic data for this case control dataset was supported by the National Institute for Health Research, Biomedical Research Centre at University College London Hospital. **ALSPAC-M** phenotypic data was collected with funding from the British Heart Foundation (SP/07/008/24066), Wellcome Trust (WT092830M) and UK Research Councils (UKRC) via the MRC (G1001357); genetic data collection was funded by the Wellcome Trust (WT088806). In addition the ALSPAC full study receives core support from The University of Bristol, UK Medical Research Council and the Wellcome Trust (102215/2/13/2) and the University of Bristol. The ALSPAC team is extremely grateful to all the families who took part in this study, the midwives for their help in recruiting them, and the whole ALSPAC team, which includes interviewers, computer and laboratory technicians, clerical workers, research scientists, volunteers, managers, receptionists and nurses.

Summary genome-wide association data on adiponectin have been contributed by ADIPOGen Consortium and have been downloaded from https://www.mcgill.ca/genepi/adipogen-consortium. Summary genome-wide association data on metabolic measures have been contributed by Kettunen et al. (30) and have been downloaded from https://www.mcgill.ca/genepi/adipogen-consortium.

## SUPPLEMENTARY METHODS

### Mendelian randomization analyses

The two-sample Mendelian randomization estimates and respective standard errors were obtained using the inverse variance-weighted (IVW) method with the following formulas:

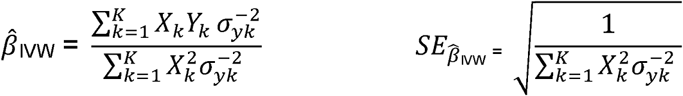

Where X_k_ is the mean change in standardized log adiponectin units per additional effect allele of SNP k and Y_k_ is the mean change in standardized units of metabolic measures per additional effect allele of SNP k with standard error σ_Yk_. To increase precision and avoid bias due to statistical overfitting, estimates for X_k_ were obtained from ADIPOGen consortium dataset (42). Prior to analysis, estimates from ADIPOGen consortium were standardized (converted from log adiponectin to standardized log adiponectin units) using individual level data from PEL82 with a similar adiponectin distribution (adiponectin concentration in ADIPOGen consortium: mean = 9.8 µg/ml (standard deviation = 5.6); adiponectin concentration in 1982 Pelotas Birth Cohort: mean = 9.3 µg/ml (standard deviation = 5.7)). Estimates for Y_k_ were derived from each study using linear regression models considering an additive model for SNP alleles.

### Comparison between multivariable and Mendelian randomization analyses

Results from conventional multivariable and Mendelian randomization analyses were compared using the Z-test:

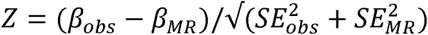

Where β_obs_ represents estimates from conventional observational analysis (with respective standard error, SE_obs_) and β_MR_ represents estimates from Mendelian randomization analysis (with respective standard error, SE_MR_).

### Proportion of variance in adiponectin concentration explained by genetic instruments

In order to estimate the strength of our genetic instruments, we estimated the phenotypic variance explained by a given SNP (R^2^) for adiponectin concentration. We used ADIPOGen summary data to approximate R^2^ for a given SNP based on the effect estimate for its association with the trait of interest (beta or 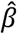), respective standard error (*se*(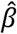)), minor allele frequency (MAF), and sample size (N). The following formula was used as previously described by Shim et al., 2015 (52):

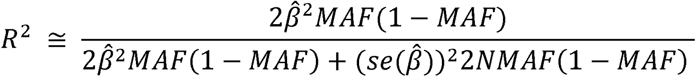

The phenotypic variance explained by the composite genetic instrument (combining all SNPs) was estimated by the sum of SNP-specific R^2^ as shown below:

SNPs used as instrumental variables for adiponectin concentration in Mendelian randomization analysis and association with adiponectin concentration

| rs ID | Chr | EA | NEA | EAF | $R^2$ | N |
| --- | --- | --- | --- | --- | --- | --- |
| rs6810075 | 3 | T | C | 0.63 | 0.0066 | 29140 |
| rs16861209 | 3 | A | C | 0.01 | 0.0125 | 29199 |
| rs17366568 | 3 | G | A | 0.91 | 0.0125 | 24865 |
| rs3774261 | 3 | A | G | 0.60 | 0.0080 | 29081 |
| Combined instrument | N/A | N/A | N/A | N/A | 0.0396 | N/A |

Chr: chromosome; EA: effect allele (trait-increasing allele); NEA: non-effect allele; $R^2$ : proportion of phenotypic variance explained by SNP; Beta: increase in standardized log adiponectin concentration per EA; SE: standard error; N: sample size; N/A: non applicable.

**Supplementary figure 1.**
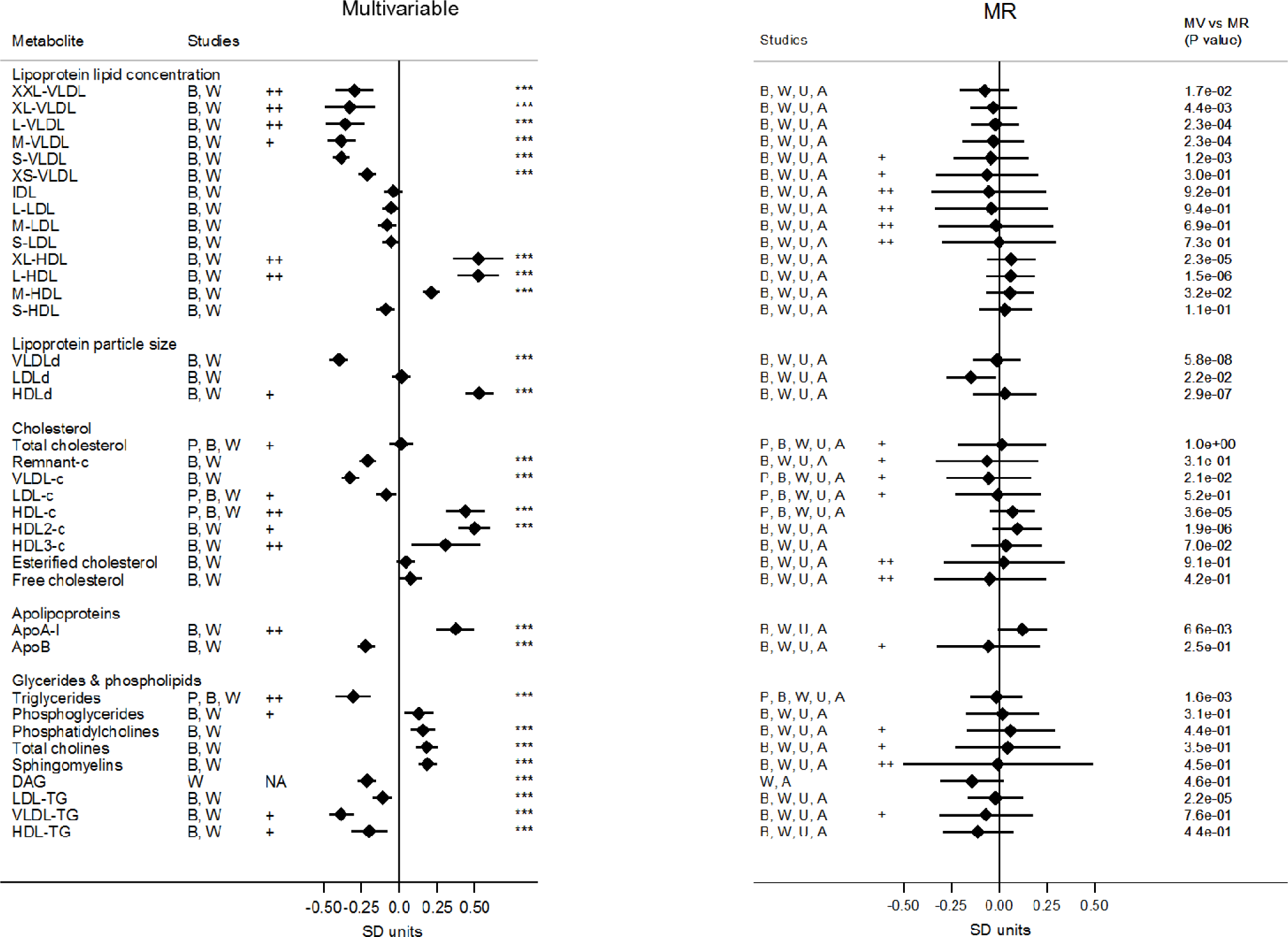
Association of lipoprotein traits with blood adiponectin levels from observational and Mendelian randomization (MR) analysis among women. Values are expressed as units of standardized log metabolite concentration (and 95% CI) per 1 unit increment of standardized log adiponectin levels. P-values for the association between adiponectin and metabolites are indicated by three asterisks (“***”) if lower than Bonferroni-adjusted threshold (P-value < 0.00068). Heterogeneity was considered substantial if I^2^ = 50-75% (“+”) or very high if I^2^ > 75% (“++”). P-values for the comparison between multivariable and Mendelian randomization estimates are displayed in the column “MR vs MV (P-value)”. Metabolic measures were adjusted for age, sex, and, if applicable, place of recruitment (BWHHS and UKCTOCS) or principal components of genomic ancestry (PEL82 and some studies contributing to Metabolomics consortium) and the resulting residuals were transformed to normal distribution by inverse rank-based normal transformation. XXL: extremely large, XL: very large, L: large, M: medium, S: small, XS: very small, VLDL: very low-density lipoprotein, LDL: low-density lipoprotein, IDL: intermediate-density lipoprotein, HDL: high-density lipoprotein, c: cholesterol, DAG: diglycerides, TG: triglycerides, P: 1982 Pelotas Birth Cohort, B: British Women Heart and Health Study, W: Whitehall II Study, U: UKCTOCS nested case-control study, A: The Avon Longitudinal Study of Children and Parents – mothers’ cohort, SD units: standard deviation units, CI: confidence interval.

**Supplementary figure 2.**
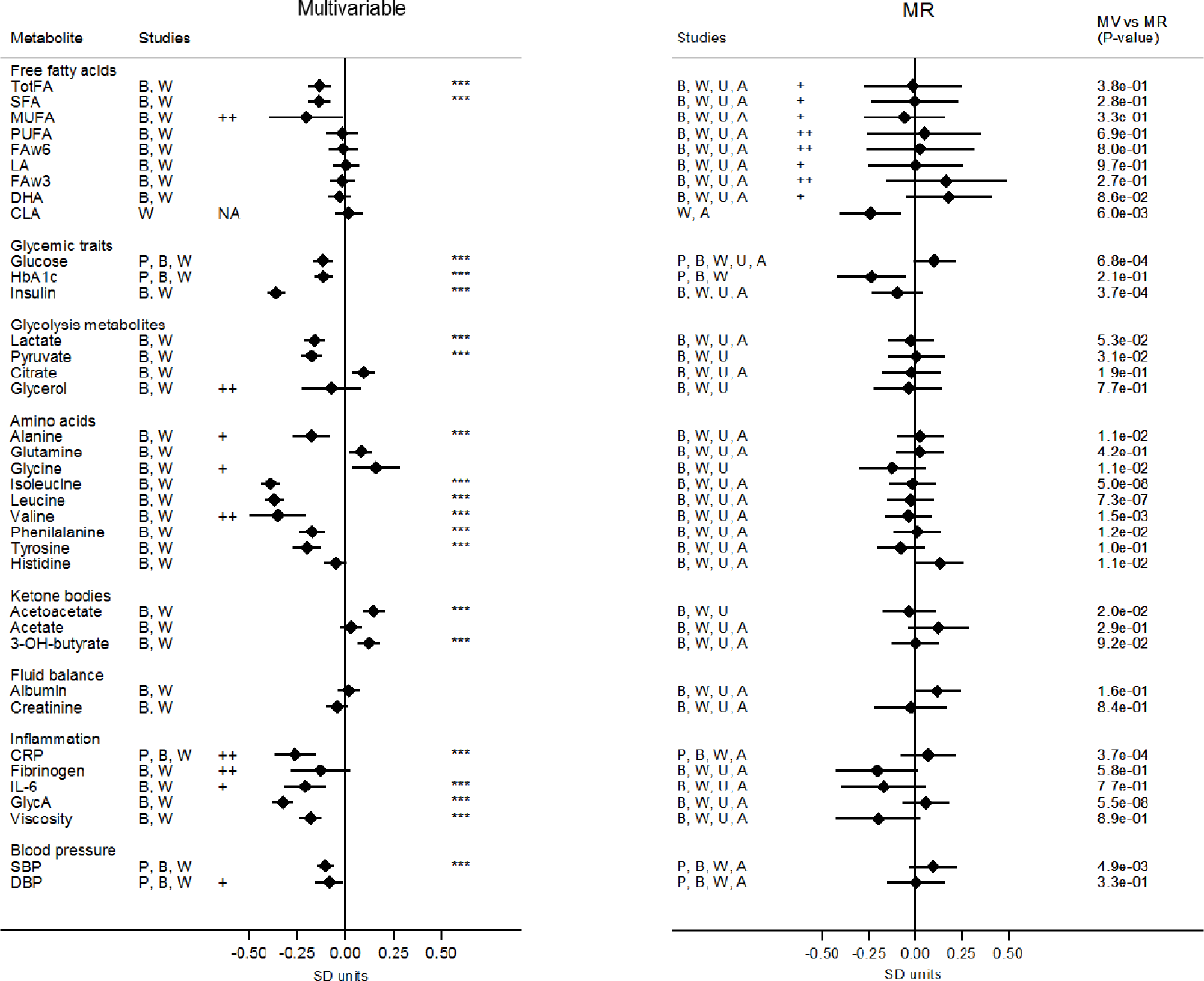
Association of multiple metabolic measures with blood adiponectin levels from observational and Mendelian randomization analysis among women. Values are expressed as units of standardized log metabolite concentration (and 95% CI) per 1 unit increment of standardized log adiponectin levels. P-values for the association between adiponectin and metabolites are indicated by three asterisks (“***”) if lower than Bonferroni-adjusted threshold (P-value < 0.00068). Heterogeneity was considered substantial if I^2^ = 50-75% (“+”) or very high if I^2^ > 75% (“++”). P-values for the comparison between multivariable and Mendelian randomization estimates are displayed in the column “MR vs MV (P-value)”. Metabolic measures were adjusted for age, sex, and, if applicable, place of recruitment (BWHHS and UKCTOCS) or principal components of genomic ancestry (PEL82 and some studies contributing to Metabolomics consortium) and the resulting residuals were transformed to normal distribution by inverse rank-based normal transformation. TotFA: total fatty acids, SFA: saturated fatty acid, MUFA: monounsaturated fatty acid, PUFA: polyunsaturated fatty acids, FAw6: omega-6 fatty acid, LA: linoleic acid, FAw3: omega-3 fatty acid, DHA: docosaexaenoic acid, CLA: conjugated linoleic acids, HbA1c: glycated haemoglobin, CRP: c-reactive protein, IL-6: interleukin-6, GlycA: glycoprotein acetyls, SBP: systolic blood pressure, DBP: diastolic blood pressure, P: 1982 Pelotas Birth Cohort, B: British Women Heart and Health Study, W: Whitehall II Study, U: UKCTOCS nested case-control study, A: The Avon Longitudinal Study of Children and Parents – mothers’ cohort, SD units: standard deviation units, CI: confidence interval.

**Supplementary figure 3.**
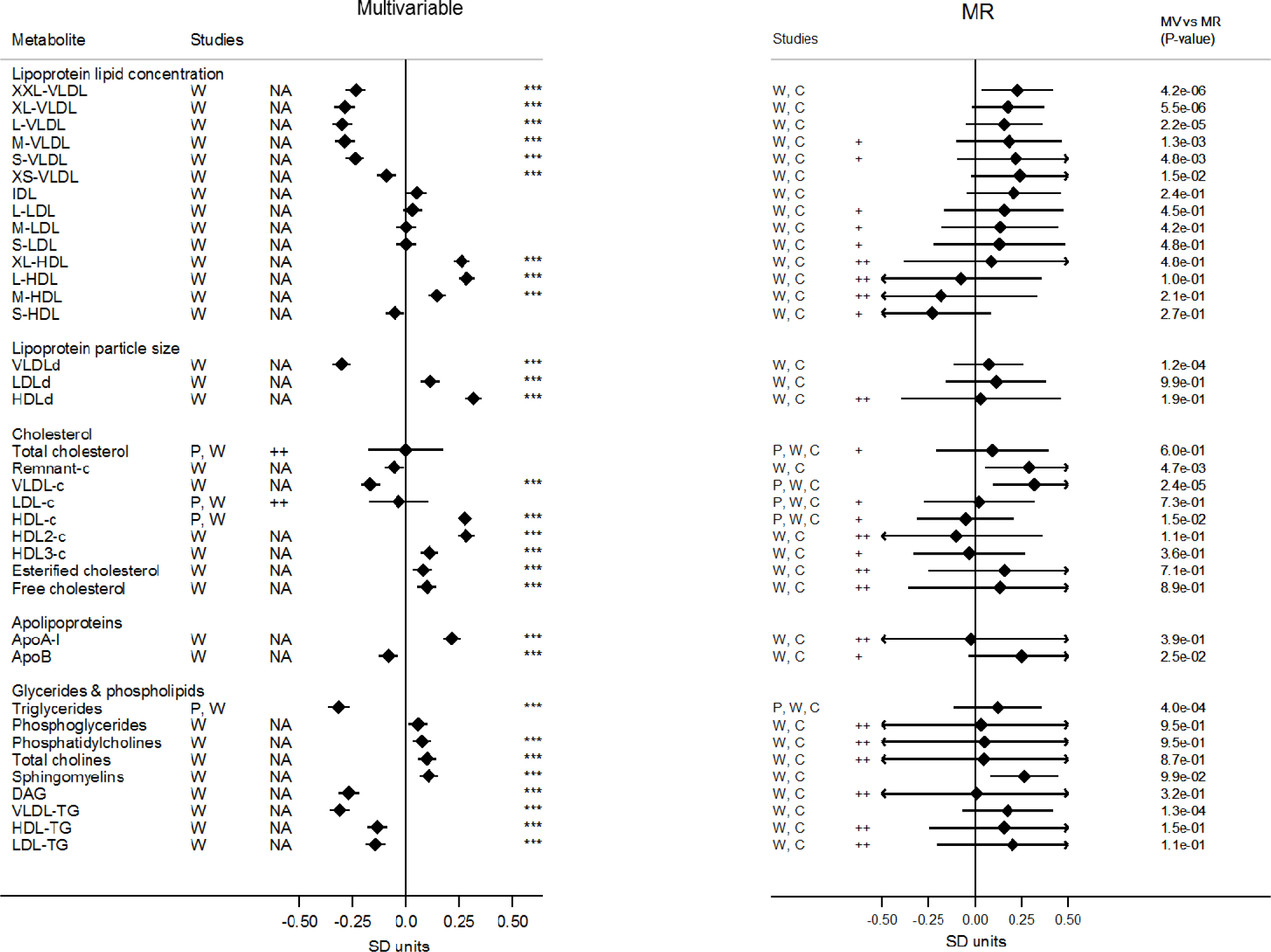
Association of lipoprotein traits with blood adiponectin levels from observational and Mendelian randomization (MR) analysis among men. Values are expressed as units of standardized log metabolite concentration (and 95% CI) per 1 unit increment of standardized log adiponectin levels. P-values for the association between adiponectin and metabolites are indicated by three asterisks (“***”) if lower than Bonferroni-adjusted threshold (P-value < 0.00068). Heterogeneity was considered substantial if I^2^ = 50-75% (“+”), very high if I^2^ > 75% (“++”) or not applicable (“NA”) when only one study contributed to the estimate. P-values for the comparison between multivariable and Mendelian randomization estimates are displayed in the column “MR vs MV (P-value)”. Metabolic measures were adjusted for age, sex, and, if applicable, place of recruitment (BWHHS and UKCTOCS) or principal components of genomic ancestry (PEL82 and some studies contributing to Metabolomics consortium) and the resulting residuals were transformed to normal distribution by inverse rank-based normal transformation. XXL: extremely large, XL: very large, L: large, M: medium, S: small, XS: very small, VLDL: very low-density lipoprotein, LDL: low-density lipoprotein, IDL: intermediate-density lipoprotein, HDL: high-density lipoprotein, c: cholesterol, DAG: diglycerides, TG: triglycerides, P: 1982 Pelotas Birth Cohort, W: Whitehall II Study, C: The Caerphilly Prospective Study, SD units: standard deviation units, CI: confidence interval.

**Supplementary figure 4.**
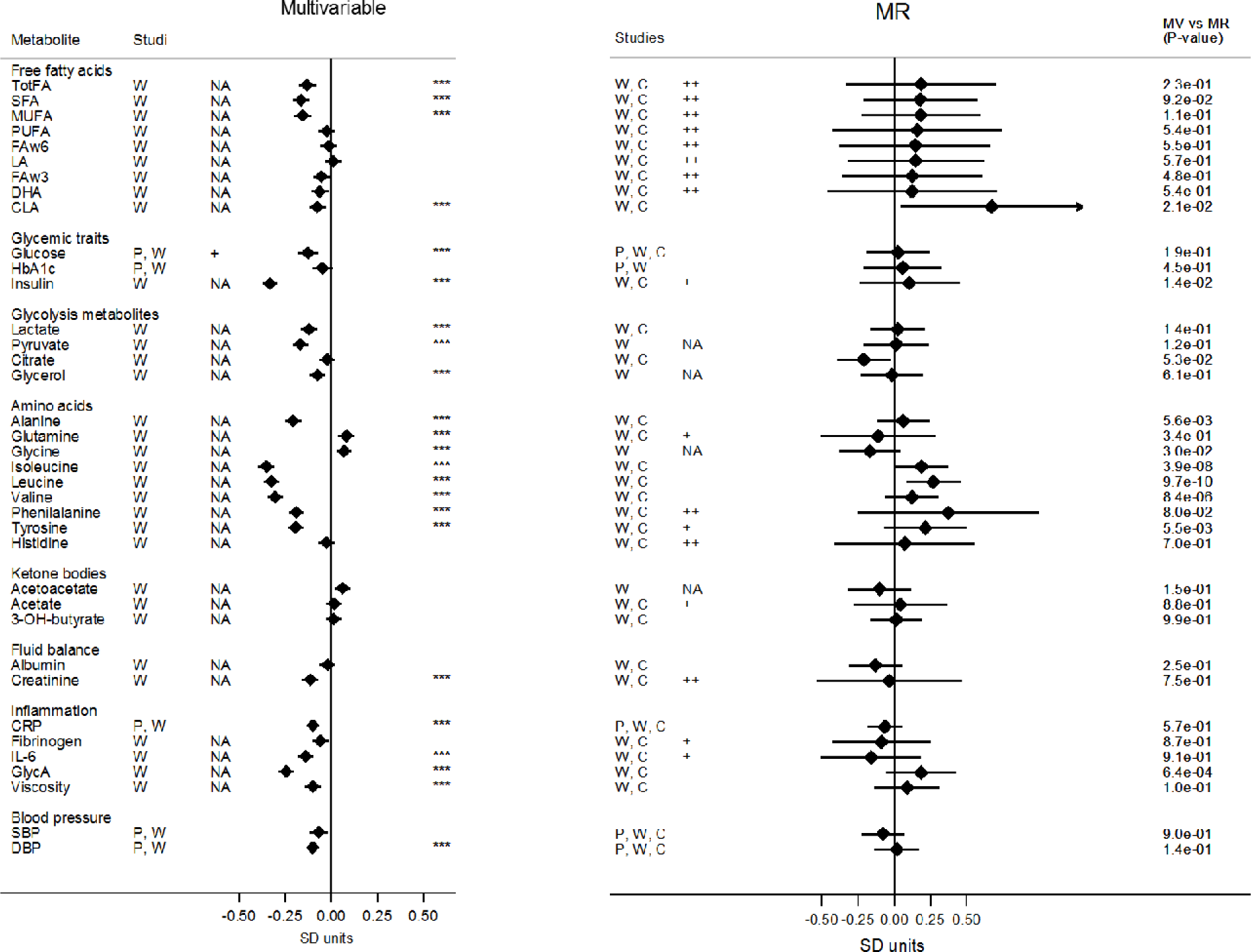
Association of multiple metabolic measures with blood adiponectin levels from observational and Mendelian randomization analysis among men. Values are expressed as units of standardized log metabolite concentration (and 95% CI) per 1 unit increment of standardized log adiponectin levels. P-values for the association between adiponectin and metabolites are indicated by three asterisks (“***”) if lower than Bonferroni-adjusted threshold (P-value < 0.00068). Heterogeneity was considered substantial if I^2^ = 50-75% (“+”) or very high if I^2^ > 75% (“++”) or not applicable (“NA”) when only one study contributed to the estimate. P-values for the comparison between multivariable and Mendelian randomization estimates are displayed in the column “MR vs MV (P-value)”. Metabolic measures were adjusted for age, sex, and, if applicable, place of recruitment (BWHHS and UKCTOCS) or principal components of genomic ancestry (PEL82 and some studies contributing to Metabolomics consortium) and the resulting residuals were transformed to normal distribution by inverse rank-based normal transformation. TotFA: total fatty acids, SFA: saturated fatty acid, MUFA: monounsaturated fatty acid, PUFA: polyunsaturated fatty acids, FAw6: omega-6 fatty acid, LA: linoleic acid, FAw3: omega-3 fatty acid, DHA: docosaexaenoic acid, CLA: conjugated linoleic acids, HbA1c: glycated haemoglobin, CRP: c-reactive protein, IL-6: interleukin-6, GlycA: glycoprotein acetyls, SBP: systolic blood pressure, DBP: diastolic blood pressure, P: 1982 Pelotas Birth Cohort, W: Whitehall II Study, C: The Caerphilly Prospective Study, SD units: standard deviation units, CI: confidence interval.

**Supplementary figure 5.**
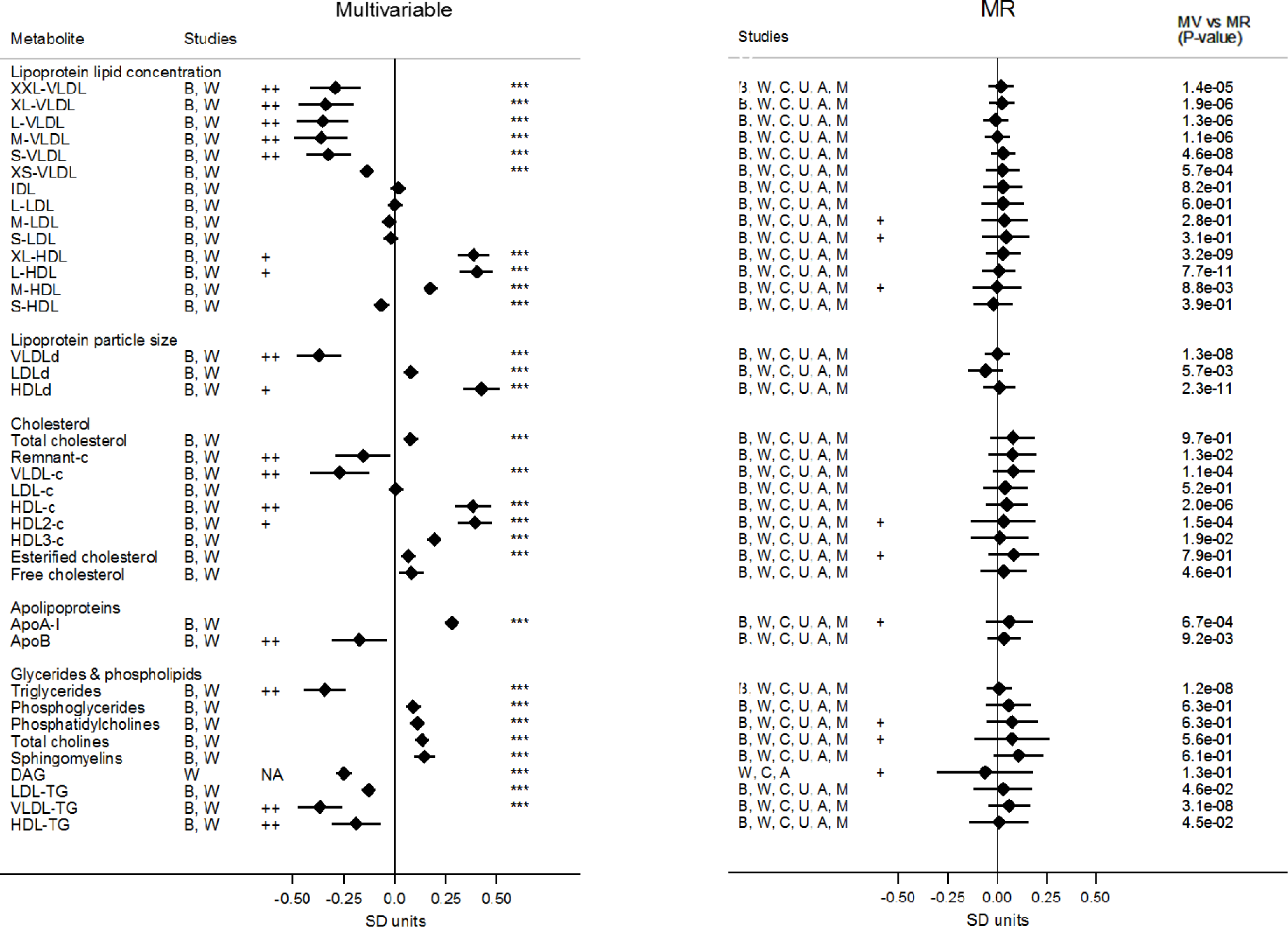
Association of lipoprotein traits with blood adiponectin levels from observational and Mendelian randomization (MR) analysis restricted to individuals of European ancestry. Values are expressed as units of standardized log metabolite concentration (and 95% CI) per 1 unit increment of standardized log adiponectin levels. P-values for the association between adiponectin and metabolites are indicated by three asterisks (“***”) if lower than Bonferroni-adjusted threshold (P-value < 0.00068). Heterogeneity was considered substantial if I^2^ = 50-75% (“+”) or very high if I^2^ > 75% (“++”). P-values for the comparison between multivariable and Mendelian randomization estimates are displayed in the column “MR vs MV (P-value)”. Metabolic measures were adjusted for age, sex, and, if applicable, place of recruitment (BWHHS and UKCTOCS) or principal components of genomic ancestry (PEL82 and some studies contributing to Metabolomics consortium) and the resulting residuals were transformed to normal distribution by inverse rank-based normal transformation. XXL: extremely large, XL: very large, L: large, M: medium, S: small, XS: very small, VLDL: very low-density lipoprotein, LDL: low-density lipoprotein, IDL: intermediate-density lipoprotein, HDL: high-density lipoprotein, c: cholesterol, DAG: diglycerides, TG: triglycerides, P: 1982 Pelotas Birth Cohort, B: British Women Heart and Health Study, W: Whitehall II Study, U: UKCTOCS nested case-control study, A: The Avon Longitudinal Study of Children and Parents – mothers’ cohort, M: Metabolomics consortium, SD units: standard deviation units, CI: confidence interval.

**Supplementary figure 6.**
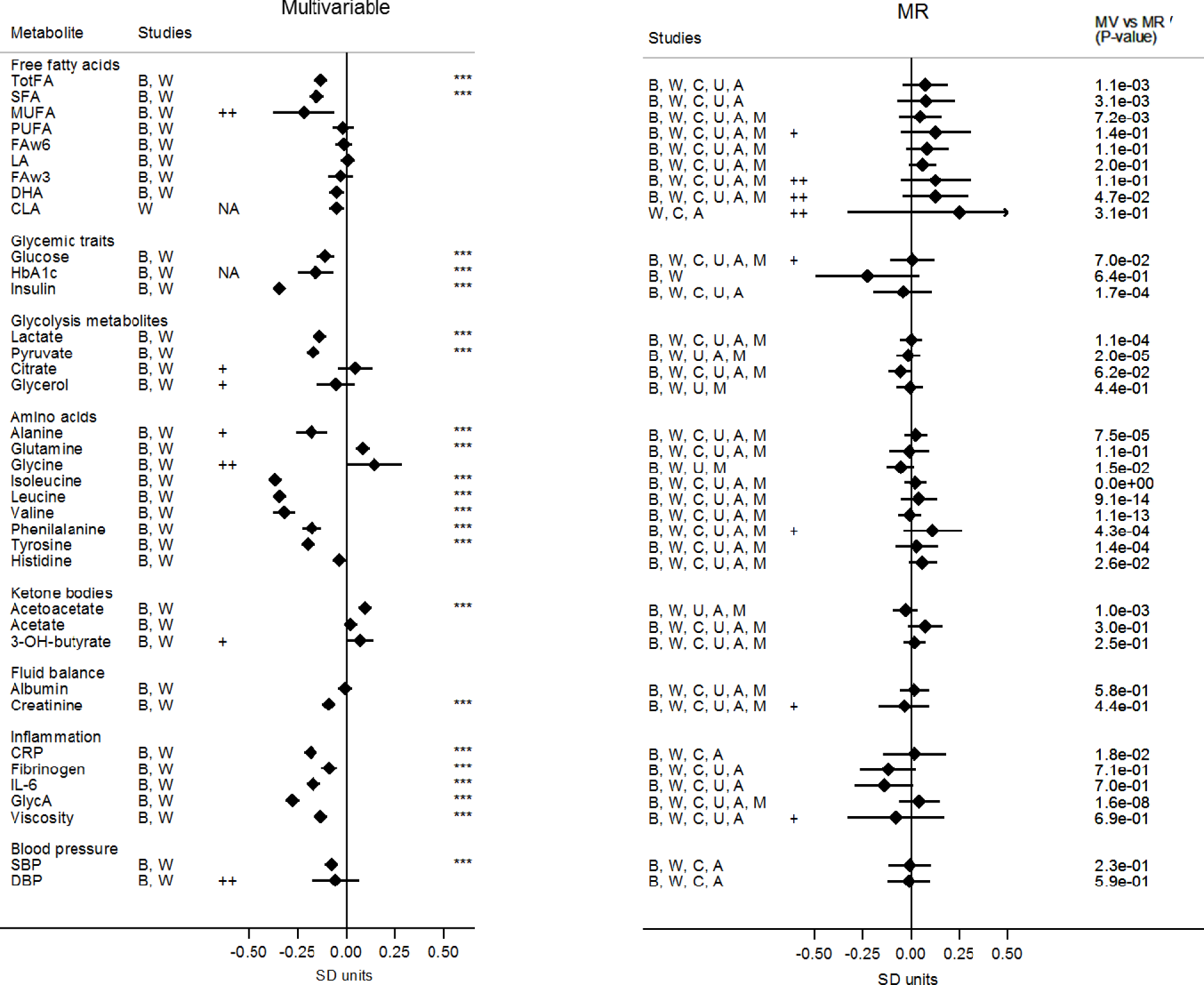
Association of multiple metabolic measures with blood adiponectin levels from observational and Mendelian randomization analysis restricted to individuals of European ancestry. Values are expressed as units of standardized log metabolite concentration (and 95% CI) per 1 unit increment of standardized log adiponectin levels. P-values for the association between adiponectin and metabolites are indicated by three asterisks (“***”) if lower than Bonferroni-adjusted threshold (P-value < 0.00068). Heterogeneity was considered substantial if I^2^ = 50-75% (“+”) or very high if I^2^ > 75% (“++”). P-values for the comparison between multivariable and Mendelian randomization estimates are displayed in the column “MR vs MV (P-value)”. Metabolic measures were adjusted for age, sex, and, if applicable, place of recruitment (BWHHS and UKCTOCS) or principal components of genomic ancestry (PEL82 and some studies contributing to Metabolomics consortium) and the resulting residuals were transformed to normal distribution by inverse rank-based normal transformation. TotFA: total fatty acids, SFA: saturated fatty acid, MUFA: monounsaturated fatty acid, PUFA: polyunsaturated fatty acids, FAw6: omega-6 fatty acid, LA: linoleic acid, FAw3: omega-3 fatty acid, DHA: docosaexaenoic acid, CLA: conjugated linoleic acids, HbA1c: glycated haemoglobin, CRP: c-reactive protein, IL-6: interleukin-6, GlycA: glycoprotein acetyls, SBP: systolic blood pressure, DBP: diastolic blood pressure, P: 1982 Pelotas Birth Cohort, B: British Women Heart and Health Study, W: Whitehall II Study, U: UKCTOCS nested case-control study, A: The Avon Longitudinal Study of Children and Parents – mothers’ cohort, M: Metabolomics consortium, SD units: standard deviation units, CI: confidence interval.

**Supplementary figure 7.**
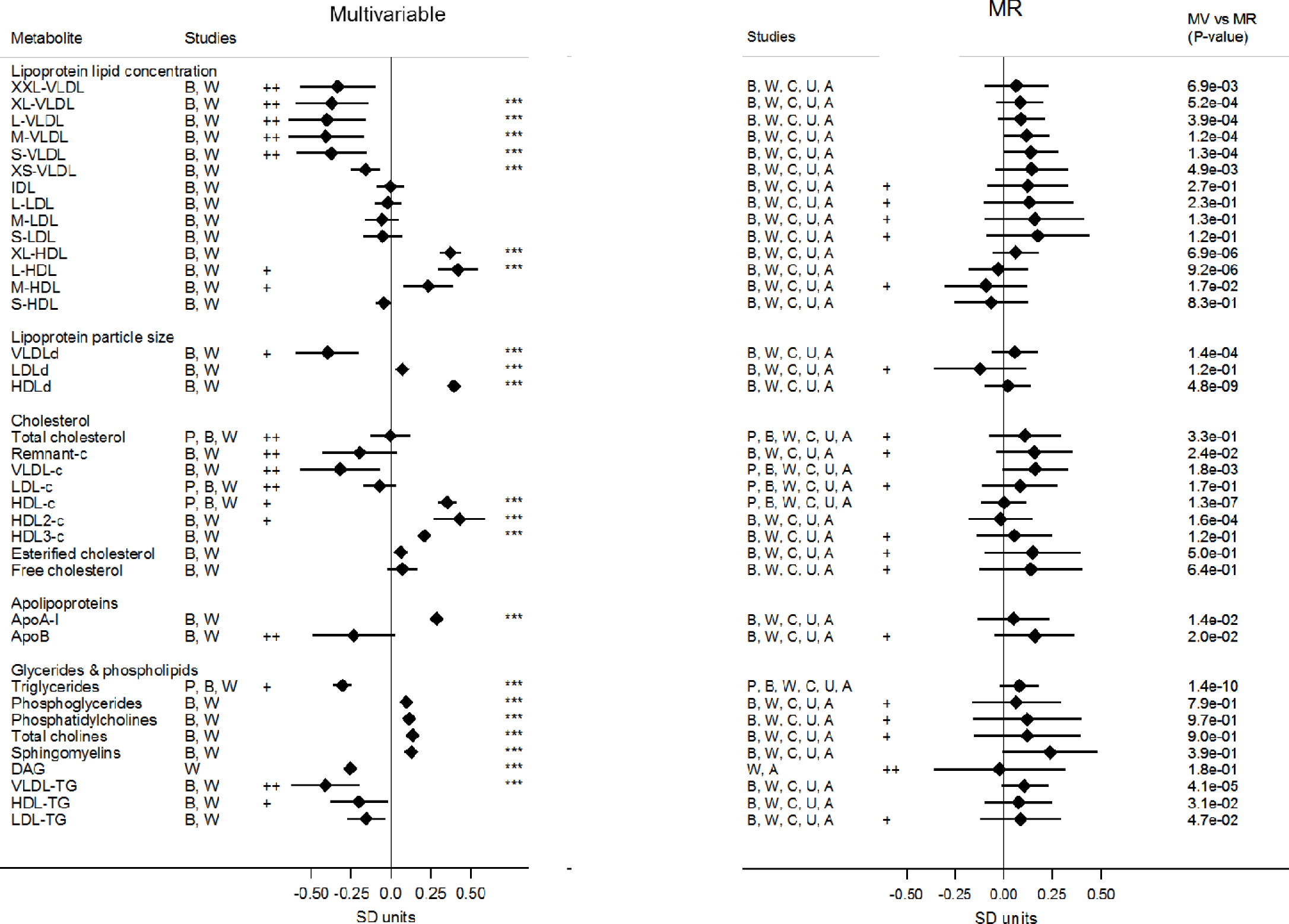
Association of lipoprotein traits with blood adiponectin levels from observational and Mendelian randomization (MR) analysis among younger individuals (< 65 years old) free from cardiovascular disease. Values are expressed as units of standardized log metabolite concentration (and 95% CI) per 1 unit increment of standardized log adiponectin levels. P-values for the association between adiponectin and metabolites are indicated by three asterisks (“***”) if lower than Bonferroni-adjusted threshold (P-value < 0.00068). Heterogeneity was considered substantial if I^2^ = 50-75% (“+”) or very high if I^2^ > 75% (“++”). P-values for the comparison between multivariable and Mendelian randomization estimates are displayed in the column “MR vs MV (P-value)”. Metabolic measures were adjusted for age, sex, and, if applicable, place of recruitment (BWHHS and UKCTOCS) or principal components of genomic ancestry (PEL82 and some studies contributing to Metabolomics consortium) and the resulting residuals were transformed to normal distribution by inverse rank-based normal transformation. XXL: extremely large, XL: very large, L: large, M: medium, S: small, XS: very small, VLDL: very low-density lipoprotein, LDL: low-density lipoprotein, IDL: intermediate-density lipoprotein, HDL: high-density lipoprotein, c: cholesterol, DAG: diglycerides, TG: triglycerides, P: 1982 Pelotas Birth Cohort, B: British Women Heart and Health Study, W: Whitehall II Study, U: UKCTOCS nested case-control study, A: The Avon Longitudinal Study of Children and Parents – mothers’ cohort, SD units: standard deviation units, CI: confidence interval.

**Supplementary figure 8.**
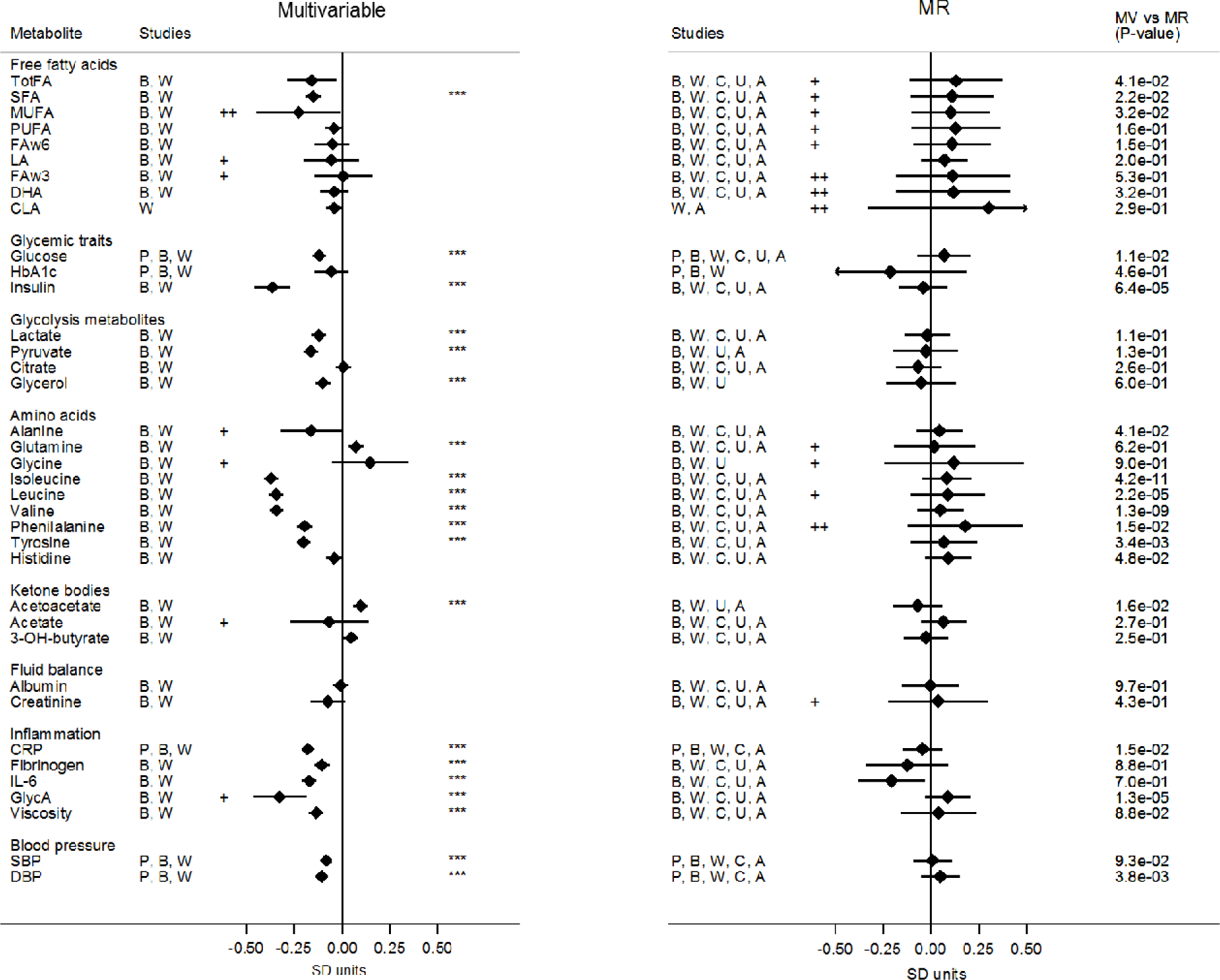
Association of multiple metabolic measures with blood adiponectin levels from observational and Mendelian randomization analysis among younger individuals (< 65 years old) free from cardiovascular disease. Values are expressed as units of standardized log metabolite concentration (and 95% CI) per 1 unit increment of standardized log adiponectin levels. P-values for the association between adiponectin and metabolites are indicated by three asterisks (“***”) if lower than Bonferroni-adjusted threshold (P-value < 0.00068). Heterogeneity was considered substantial if I^2^ = 50-75% (“+”) or very high if I^2^ > 75% (“++”). P-values for the comparison between multivariable and Mendelian randomization estimates are displayed in the column “MR vs MV (P-value)”. Metabolic measures were adjusted for age, sex, and, if applicable, place of recruitment (BWHHS and UKCTOCS) or principal components of genomic ancestry (PEL82 and some studies contributing to Metabolomics consortium) and the resulting residuals were transformed to normal distribution by inverse rank-based normal transformation. TotFA: total fatty acids, SFA: saturated fatty acid, MUFA: monounsaturated fatty acid, PUFA: polyunsaturated fatty acids, FAw6: omega-6 fatty acid, LA: linoleic acid, FAw3: omega-3 fatty acid, DHA: docosaexaenoic acid, CLA: conjugated linoleic acids, HbA1c: glycated haemoglobin, CRP: c-reactive protein, IL-6: interleukin-6, GlycA: glycoprotein acetyls, SBP: systolic blood pressure, DBP: diastolic blood pressure, P: 1982 Pelotas Birth Cohort, B: British Women Heart and Health Study, W: Whitehall II Study, U: UKCTOCS nested case-control study, A: The Avon Longitudinal Study of Children and Parents – mothers’ cohort, M: Metabolomics consortium, SD units: standard deviation units, CI: confidence interval.

**Supplementary table 1.**
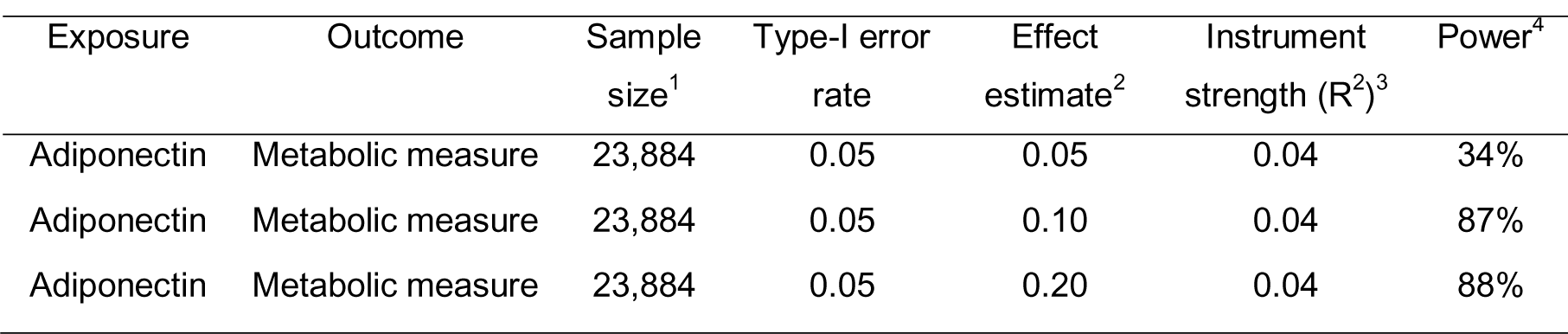
Estimated power in Mendelian randomization analysis.

**Supplementary table 2.** Heterogeneity estimates (I^2^) for meta-analysis of study-specific multivariate (MV) and Mendelian randomization (MR) estimates.

| Metabolite | Overall |  | Females |  | Males |  | European studies only |  | Low risk individuals only |  |
| --- | --- | --- | --- | --- | --- | --- | --- | --- | --- | --- |
|  | MV | MR | MV | MR | MV | MR | MV | MR | MV | MR |
| XXL-VLDL | 85 | 0 | 82 | 7 | NA | 0 | 85 | 0 | 78 | 34 |
| XL-VLDL | 88 | 0 | 91 | 0 | NA | 0 | 88 | 0 | 79 | 0 |
| L-VLDL | 86 | 0 | 84 | 0 | NA | 11 | 86 | 0 | 82 | 0 |
| M-VLDL | 88 | 0 | 68 | 35 | NA | 52 | 88 | 0 | 83 | 0 |
| S-VLDL | 81 | 0 | 0 | 52 | NA | 63 | 81 | 0 | 79 | 19 |
| XS-VLDL | 0 | 23 | 0 | 73 | NA | 49 | 0 | 23 | 23 | 48 |
| IDL | 0 | 36 | 0 | 78 | NA | 44 | 0 | 36 | 23 | 57 |
| L-LDL | 0 | 43 | 0 | 78 | NA | 64 | 0 | 43 | 20 | 65 |
| M-LDL | 0 | 51 | 0 | 79 | NA | 63 | 0 | 51 | 32 | 71 |
| S-LDL | 0 | 53 | 0 | 78 | NA | 71 | 0 | 53 | 40 | 73 |
| XL-HDL | 66 | 26 | 88 | 0 | NA | 84 | 66 | 26 | 19 | 0 |
| L-HDL | 72 | 22 | 83 | 0 | NA | 81 | 72 | 22 | 59 | 28 |
| M-HDL | 0 | 57 | 0 | 0 | NA | 86 | 0 | 57 | 62 | 58 |
| S-HDL | 0 | 36 | 0 | 8 | NA | 63 | 0 | 36 | 4 | 50 |
| VLDLd | 83 | 0 | 26 | 0 | NA | 0 | 83 | 0 | 73 | 0 |
| LDLd | 0 | 25 | 0 | 0 | NA | 50 | 0 | 25 | 0 | 66 |
| HDLd | 75 | 18 | 64 | 32 | NA | 80 | 75 | 18 | 0 | 0 |
| Total cholesterol | 94 | 44 | 71 | 73 | 96 | 73 | 0 | 49 | 92 | 63 |
| Remnant-c | 87 | 23 | 0 | 73 | NA | 36 | 87 | 23 | 76 | 52 |
| VLDL-c | 89 | 4 | 0 | 61 | NA | 26 | 89 | 4 | 81 | 38 |
| LDL-c | 91 | 45 | 61 | 71 | 94 | 73 | 0 | 47 | 89 | 66 |
| HDL-c | 75 | 33 | 90 | 0 | 0 | 68 | 76 | 42 | 73 | 16 |
| HDL2-c | 74 | 57 | 70 | 0 | NA | 83 | 74 | 57 | 72 | 34 |
| HDL3-c | 0 | 45 | 93 | 43 | NA | 60 | 0 | 45 | 0 | 52 |
| Esterified cholesterol | 0 | 56 | 0 | 80 | NA | 77 | 0 | 56 | 0 | 68 |
| Free cholesterol | 43 | 46 | 31 | 77 | NA | 85 | 43 | 46 | 27 | 73 |
| ApoA-I | 0 | 55 | 79 | 0 | NA | 88 | 0 | 55 | 0 | 46 |
| ApoB | 86 | 20 | 0 | 73 | NA | 56 | 86 | 20 | 81 | 56 |
| Triglycerides | 73 | 0 | 91 | 39 | 48 | 50 | 80 | 0 | 70 | 0 |
| Phosphoglycerides | 0 | 48 | 58 | 48 | NA | 91 | 0 | 48 | 0 | 62 |
| Phosphatidylcholines | 0 | 58 | 41 | 62 | NA | 92 | 0 | 58 | 0 | 75 |
| Total cholines | 0 | 67 | 29 | 73 | NA | 90 | 0 | 67 | 0 | 74 |
| Sphingomyelins | 30 | 43 | 0 | 85 | NA | 0 | 30 | 43 | 0 | 45 |
| DAG | NA | 70 | NA | 0 | NA | 86 | NA | 70 | NA | 84 |
| HDL-TG | 0 | 47 | 75 | 44 | NA | 76 | 82 | 0 | 79 | 0 |
| VLDL-TG | 84 | 49 | 60 | 22 | NA | 36 | 84 | 49 | 68 | 40 |
| LDL-TG | 82 | 0 | 19 | 68 | NA | 78 | 0 | 47 | 42 | 56 |
| TotFA | 0 | 48 | 5 | 72 | NA | 86 | 0 | 48 | 42 | 68 |
| SFA | 0 | 45 | 0 | 65 | NA | 76 | 0 | 45 | 0 | 61 |
| MUFA | 91 | 43 | 91 | 61 | NA | 77 | 91 | 43 | 77 | 55 |
| PUFA | 32 | 64 | 45 | 79 | NA | 89 | 32 | 64 | 0 | 66 |
| FAw6 | 10 | 42 | 35 | 77 | NA | 86 | 10 | 42 | 22 | 54 |
| LA | 0 | 0 | 16 | 70 | NA | 83 | 0 | 0 | 53 | 3 |
| FAw3 | 47 | 79 | 13 | 82 | NA | 84 | 47 | 79 | 60 | 80 |
| DHA | 0 | 76 | 0 | 64 | NA | 89 | 0 | 76 | 16 | 80 |
| CLA | NA | 92 | NA | 0 | NA | 46 | NA | 92 | NA | 93 |
| Glucose | 0 | 54 | 46 | 0 | 57 | 49 | 21 | 51 | 34 | 39 |
| HbA1c | 62 | 0 | 11 | 0 | NA | NA | NA | NA | 21 | 38 |
| Insulin | 0 | 38 | 0 | 0 | NA | 53 | 0 | 38 | 37 | 0 |
| Lactate | 0 | 0 | 0 | 0 | NA | 0 | 0 | 0 | 0 | 0 |
| Pyruvate | 0 | 0 | 0 | 24 | NA | NA | 0 | 0 | 0 | 27 |
| Citrate | 69 | 0 | 0 | 29 | NA | 0 | 69 | 0 | 0 | 0 |
| Glycerol | 74 | 0 | 83 | 0 | NA | NA | 74 | 0 | 0 | 0 |
| Alanine | 65 | 0 | 62 | 0 | NA | 0 | 65 | 0 | 58 | 0 |
| Glutamine | 0 | 43 | 0 | 0 | NA | 75 | 0 | 43 | 0 | 58 |
| Glycine | 87 | 0 | 71 | 0 | NA | NA | 87 | 0 | 69 | 55 |
| Isoleucine | 0 | 0 | 0 | 0 | NA | 0 | 0 | 0 | 0 | 9 |
| Leucine | 0 | 34 | 0 | 0 | NA | 0 | 0 | 34 | 0 | 52 |
| Valine | 45 | 0 | 87 | 0 | NA | 0 | 45 | 0 | 0 | 0 |
| Phenylalanine | 32 | 72 | 30 | 0 | NA | 91 | 32 | 72 | 0 | 79 |
| Tyrosine | 0 | 48 | 40 | 0 | NA | 54 | 0 | 48 | 0 | 42 |
| Histidine | 0 | 10 | 0 | 0 | NA | 84 | 0 | 10 | 0 | 0 |
| Acetoacetate | 0 | 0 | 0 | 14 | NA | NA | 0 | 0 | 0 | 0 |
| Acetate | 0 | 31 | 0 | 35 | NA | 64 | 0 | 31 | 74 | 0 |
| 3-OH-butyrate | 54 | 0 | 0 | 0 | NA | 0 | 54 | 0 | 0 | 0 |
| Albumin | 0 | 11 | 0 | 0 | NA | 0 | 0 | 11 | 0 | 26 |
| Creatinine | 0 | 61 | 0 | 49 | NA | 85 | 0 | 61 | 22 | 71 |
| CRP | 0 | 28 | 81 | 13 | 41 | 0 | 0 | 45 | 0 | 0 |
| Fibrinogen | 16 | 0 | 87 | 0 | NA | 60 | 16 | 0 | 0 | 17 |
| IL-6 | 0 | 3 | 69 | 0 | NA | 57 | 0 | 3 | 0 | 0 |
| GlycA | 0 | 43 | 0 | 0 | NA | 39 | 0 | 43 | 58 | 0 |
| Viscosity | 0 | 56 | 0 | 0 | NA | NA | 0 | 56 | 0 | 0 |
| SBP | 0 | 0 | 28 | 11 | 44 | 0 | 0 | 0 | 0 | 0 |
| DBP | 68 | 0 | 72 | 36 | 0 | 0 | 83 | 0 | 0 | 0 |

NA: not applicable (estimates from only one study available). XXL: extremely large, XL: very large, L: large, M: medium, S: small, XS: very small, VLDL: very low-density lipoprotein, LDL: low-density lipoprotein, IDL: intermediate-density lipoprotein, HDL: high-density lipoprotein, c: cholesterol, DAG: diglycerides, TG: triglycerides, TotFA: total fatty acids, SFA: saturated fatty acid, MUFA: monounsaturated fatty acid, PUFA: polyunsaturated fatty acids, FAW6: omega-6 fatty acid, LA: linoleic acid, FAW3: omega-3 fatty acid, DHA: docosaenoic acid, CLA: conjugated linoleic acids, HbA1c: glycated haemoglobin, CRP: c-reactive protein, IL-6: interleukin-6, GlycA: glycoprotein acetyls, SBP: systolic blood pressure, DBP: diastolic blood pressure.

**Supplementary table 3.** P-values for the association of demographic and lifestyle variables with SNPs selected for Mendelian randomization analysis for each participating study.

|  | PEL82 | BWHHS | WHII | CaPS | UKTOCS<br>case-control* | ALSPAC-M |
| --- | --- | --- | --- | --- | --- | --- |
|  | *P-value* |  |  |  |  |  |
| *Sex (male vs female)* |  |  |  |  |  |  |
| rs6810075 | 0.15 | — | 0.67 | — | — | — |
| rs16861209 | 0.12 | — | 0.45 | — | — | — |
| rs17366568 | 0.36 | — | 0.84 | — | — | — |
| rs3774261 | 0.35 | — | 0.63 | — | — | — |
| *Age (years)* |  |  |  |  |  |  |
| rs6810075 | 0.78 | 0.75 | 0.28 | 0.36 | 0.59 | 0.001 |
| rs16861209 | 0.56 | 0.58 | 0.27 | 0.01 | 0.57 | 0.83 |
| rs17366568 | 0.22 | 0.83 | 0.47 | 0.56 | 0.58 | 0.93 |
| rs3774261 | 0.68 | 0.03 | 0.96 | 0.87 | 0.43 | 0.15 |
| *European (yes vs no)* |  |  |  |  |  |  |
| rs6810075 | 0.06 | 0.50 | 0.48 | — | 0.70 | 0.41 |
| rs16861209 | 0.75 | — | 0.12 | — | 0.16 | 0.35 |
| rs17366568 | 0.45 | 0.61 | — | — | 0.62 | 0.19 |
| rs3774261 | 0.44 | 0.95 | — | — | 0.85 | — |
| *Smoking (yes vs no)* |  |  |  |  |  |  |
| rs6810075 | 0.64 | 0.22 | 0.77 | 0.48 | — | 0.11 |
| rs16861209 | 0.57 | 0.37 | 0.87 | 0.48 | — | 0.24 |
| rs17366568 | 0.45 | 0.62 | 0.44 | 0.77 | — | 0.90 |
| rs3774261 | 0.52 | 0.08 | 0.90 | 0.37 | — | 0.92 |
| *Body mass index (kg/m<sup>2</sup>)* |  |  |  |  |  |  |
| rs6810075 | 0.63 | 0.49 | 0.49 | 0.21 | 0.05 | 0.45 |
| rs16861209 | 0.39 | 0.41 | 0.59 | 0.65 | 0.20 | 0.72 |
| rs17366568 | 0.47 | 0.73 | 0.87 | 0.11 | 0.66 | 0.32 |
| rs3774261 | 0.48 | 0.65 | 0.41 | 0.44 | 0.78 | 0.04 |

ALSPAC-M: The Avon Longitudinal Study of Children and Parents – mothers' cohort; BWHHS: British Women's Heart and Health Study; CaPS: The Caerphilly Prospective Study; PEL82: 1982 Pelotas Birth Cohort; UKCTOCS: case-control study nested in The United Kingdom Collaborative Trial of Ovarian Cancer Screening; WHII: Whitehall-II Study.

